# Genome mining of the citrus pathogen *Elsinoë fawcettii*; prediction and prioritisation of candidate effectors, cell wall degrading enzymes and secondary metabolite gene clusters

**DOI:** 10.1101/2019.12.19.882142

**Authors:** Sarah Jeffress, Kiruba Arun-Chinnappa, Ben Stodart, Niloofar Vaghefi, Yu Pei Tan, Gavin Ash

**Author notes:** Corresponding author: Gavin Ash. Email addresses: Sarah Jeffress, Kiruba Arun-Chinnappa Ben Stodart, Niloofar Vaghefi Yu Pei Tan, Gavin Ash.

## Abstract

*Elsinoë fawcettii*, a necrotrophic fungal pathogen, causes citrus scab on numerous citrus varieties around the world. Known pathotypes of *E. fawcettii* are based on host range; additionally, cryptic pathotypes have been reported and more novel pathotypes are thought to exist. *E. fawcettii* produces elsinochrome, a non-host selective toxin which contributes to virulence. However, the mechanisms involved in potential pathogen-host interactions occurring prior to the production of elsinochrome are unknown, yet the host-specificity observed among pathotypes suggests a reliance upon such mechanisms. In this study we have generated a whole genome sequencing project for *E. fawcettii,* producing an annotated draft assembly 26.01 Mb in size, with 10,080 predicted gene models and low (0.37%) coverage of transposable elements. The assembly showed evidence of AT-rich regions, potentially indicating genomic regions with increased plasticity. Using a variety of computational tools, we mined the *E. fawcettii* genome for potential virulence genes as candidates for future investigation. A total of 1,280 secreted proteins and 203 candidate effectors were predicted and compared to those of other necrotrophic (*Botrytis cinerea*, *Parastagonospora nodorum*, *Pyrenophora tritici-repentis*, *Sclerotinia sclerotiorum* and *Zymoseptoria tritici*), hemibiotrophic (*Leptosphaeria maculans*, *Magnaporthe oryzae*, *Rhynchosporium commune* and *Verticillium dahliae*) and biotrophic (*Ustilago maydis*) plant pathogens. Genomic and proteomic features of known fungal effectors were analysed and used to guide the prioritisation of 77 candidate effectors of *E. fawcettii*. Additionally, 378 carbohydrate-active enzymes were predicted and analysed for likely secretion and sequence similarity with known virulence genes. Furthermore, secondary metabolite prediction indicated nine additional genes potentially involved in the elsinochrome biosynthesis gene cluster than previously described. A further 21 secondary metabolite clusters were predicted, some with similarity to known toxin producing gene clusters. The candidate virulence genes predicted in this study provide a comprehensive resource for future experimental investigation into the pathogenesis of *E. fawcettii*.

## Introduction

*Elsinoë fawcettii* Bitancourt & Jenkins, a necrotrophic fungal species within the Ascomycota phylum (class Dothideomycetes, subclass Dothideomycetidae, order Myriangiales), is a filamentous phytopathogen which causes a necrotic disease, known as citrus scab, to the leaves and fruit of a variety of citrus crops around the world. Susceptible citrus varieties include lemon (*Citrus limon*), rough lemon (*C. jambhiri*), sour orange (*C. aurantium*), Rangpur lime (*C. limonia*), Temple and Murcott tangors (*C. sinensis* x *C. reticulata*), Satsuma mandarin (*C. unshiu*), grapefruit (*C. paradisi*), Cleopatra mandarin (*C. reshni*), clementine (*C. clementina*), yuzu (*C. junos*), kinkoji (*C. obovoidea*), pomelo (*C. grandis*) and Jiangjinsuanju (*C. sunki*) [1–9]. Numerous pathotypes of *E. fawcettii* are defined by host range, including the Florida Broad Host Range (FBHR), Florida Narrow Host Range (FNHR), Tyron’s, Lemon, Jinguel, SRGC and SM, while cryptic and novel pathotypes are also reported [1, 3, 10]. Only the Tyron’s pathotype (which infects Eureka lemon, Rough lemon, clementine, Rangpur lime and Cleopatra mandarin) and the Lemon pathotype (which only infects Eureka lemon, Rough lemon, Rangpur lime) have been described in Australia [2, 3, 7], however *E. fawcettii* has reportedly been isolated from kumquat (*Fortunella* sp.), tea plant (*Camellia sinensis*) and mango (*Mangifera indica*) [11], indicating a wider range of pathotypes to be present in Australia. Additional species of *Elsinoë* found causing disease in Australia include *E. ampelina,* which causes anthracnose to grapes [12] and two *E. australis* pathotypes; one which causes scab disease to jojoba (*Simmondsia chinensis*) [13] and a second found on rare occasions on finger lime (*C. australasica*) in Queensland forest areas [14]. Species of *Elsinoë* causing crop disease in countries neighbouring Australia include *E. batatas*, which causes large yield losses in sweet potato crops in Papua New Guinea [15, 16] and *E. pyri*, which infects apples in organic orchards in New Zealand [17]. Around the world there are reportedly 75 *Elsinoë* species, the majority of which appear to be host specific [18]. While citrus scab is not thought to affect yield, it reduces the value of affected fruit on the fresh market. Australia is known for producing high quality citrus fruits for local consumption and export, and so understandably, there is great interest in protecting this valuable commodity from disease.

*E. fawcettii* is commonly described as an anamorph, reproducing asexually. Hyaline and spindle shaped conidia are produced from the centre of necrotic citrus scab lesions [19, 20]. Conidia are dispersed by water splash, requiring temperatures between 23.5-27 °C with four hours of water contact for effective host infection. Therefore, disease is favoured by warm weather with overhead watering systems or rain [21]. Only young plant tissues are vulnerable to infection; leaves are susceptible from first shoots through to half expanded and similarly fruit for 6 to 8 weeks after petal fall, while mature plants are resistant to disease [19]. Cuticle, epidermal cells and mesophyll tissue are degraded within 1 to 2 days of inoculation, hyphal colonisation proceeds and within 3 to 4 days symptoms are visible [20, 22]. After formation of necrotic scab lesions on fruit, twigs and leaves, conidia are produced from the scab pustules providing inoculum for further spread. Within 5 days, host cell walls become lignified separating infected regions from healthy cells, which is thought to limit internal spread of the pathogen [20]. The necrosis that occurs during infection is produced in response to elsinochrome, a well-known secondary metabolite (SM) of species of *Elsinoë*. Elsinochromes are red or orange pigments which can be produced in culture [23, 24]. In aerobic and light-activated conditions, reactive oxygen species are produced in response to elsinochromes in a non-host selective manner, generating an environment of cellular toxicity [25]. Elsinochrome production is required for full virulence of *E. fawcettii*, specifically the *EfPKS1* and *TSF1* genes are vital within the elsinochrome gene cluster [26, 27]. However, two points indicate that *E. fawcettii* pathogenesis is more complex than simply the result of necrotic toxin production: (I) the production of elsinochrome appears to be variable and does not correlate with virulence [28]; and (II) elsinochrome is a non-host selective toxin, yet *Elsinoë* species and *E. fawcettii* pathotypes cause disease in a host specific manner. Host-specific virulence factors targeted for interaction with distinct host proteins to overcome immune defences, prior to elsinochrome production, could explain the observed host specificity. Candidate virulence genes may include effectors and cell wall degrading enzymes. Effectors are secreted pathogen proteins, targeted to either the host cytoplasm or apoplast, which enable the pathogen to evade recognition receptor activities of the host’s defence system and, if successful, infection proceeds. Resistant hosts, however, recognise pathogen effectors using resistant (R) genes which elicit plant effector-triggered immunity and pathogenesis is unsuccessful [29, 30]. While it was previously thought that necrotrophic fungal pathogens would use only a repertoire of carbohydrate-active enzymes (CAZymes) or SM’s to infect host plants [31], there is increased awareness of their utilisation of secreted protein effectors [32–37], highlighting the importance of protein effector identification in all fungal pathogens. Frequently shared features of effectors include; a signal peptide at the N-terminal and no transmembrane helices or glycosylphosphatidylinositol (GPI) anchors. Other features less frequently shared include; small size, cysteine rich, amino acid polymorphism, repetitive regions, gene duplication, no conserved protein domains, coding sequence found nearby to transposable elements, and absence in non-pathogenic strains [38–45]. Furthermore, some appear to be unique to a species for example the necrosis-inducing protein effectors NIP1, NIP2 and NIP3 of *Rhynchosporium commune* [46] and three avirulence effectors AvrLm1, AvrLm6 and AvrLm4-7 of *Leptosphaeria maculans* [47]. Others have orthologous genes or similar domains in numerous species for example the chorismate mutase effector, Cmu1, of *Ustilago maydis* [48] and the cell death-inducing effector, MoCDIP4, of *Magnaporthe oryzae* [49]. Understandably, with such a large variety of potential features, effector identification remains challenging. Effectors are found in biotrophs, for example *U. maydis* [50–53], hemibiotrophs, such as *L. maculans* [54–56], *M. oryzae* [57, 58], *R. commune* [46] and *Verticillium dahliae* [59–61], necrotrophs, for example *Botrytis cinerea* [62, 63], *Parastagonospora nodorum* [34, 42, 64], *Pyrenophora tritici-repentis* [65], *Sclerotinia sclerotiorum* [32] and also the hemibiotroph/latent necrotroph *Zymoseptoria tritici* [66].

Genomic location has potential to be an identifying feature of virulence genes in some species, for example pathogenicity-related genes of *L. maculans*, including those coding for secreted proteins and genes potentially involved in SM biosynthesis, are found at higher rates in AT-rich genomic regions in comparison to GC-equilibrated blocks [47]. It is thought that effectors and their target host proteins co-evolve, in a constant arms race [67], presenting genomic regions with higher levels of plasticity as potential niches which harbour effector genes.

Another group of virulence factors likely to play a role in *E. fawcettii* pathogenesis are cell wall degrading enzymes (CWDE), these are CAZymes, including glycoside hydrolases, polysaccharide lyases and carbohydrate esterases, which can be secreted from fungal pathogens and promote cleavage of plant cell wall components [68–70]. Cell wall components, such as cellulose, hemicelluloses (xyloglucan and arabinoxylan) and pectin (rhamnogalacturonan I, homogalacturonan, xylogalacturonan, arabinan and rhamnogalacturonan II) [71], are targets for pathogens to degrade for nutrients and/or to overcome the physical barrier to their host. CWDE’s can include polygalacturonases, pectate lyases, and pectinesterases which promote pectin degradation [72–78], glucanases (also known as cellulase) which breaks links between glucose residues [79] and xylanases which cleave links in the xylosyl backbone of xyloglucan [80–82].

*E. fawcettii* effectors and/or CWDE’s which interact with certain host plant cell wall components could explain the observed host specificity of pathotypes. Computational prediction of genes coding for such virulence factors can lead to many candidate effectors (CE) and potential CWDE’s, leading to an overabundance of candidates which require prioritisation. This study aimed to generate an assembly of the *E. fawcettii* isolate, BRIP 53147a, through whole genome shotgun (WGS) sequencing, to identify candidate virulence genes and appropriately shortlist these predictions to improve the focus of future experimental validation procedures. Computational methods involving genomic, proteomic and comparative analyses enabled the prediction and prioritisation of CE’s and CWDE’s which may be interacting with the host plant and overcoming immune defences prior to the biosynthesis of elsinochrome. Additional genes potentially involved in the elsinochrome gene cluster were also predicted, as were additional SM clusters which may be impacting virulence of *E. fawcettii*.

## Materials and Methods

### Sequencing, assembly, gene prediction, annotation and genomic analyses

*E. fawcettii* (BRIP 53147a), collected from *C. limon* (L.) Burm.f. in Montville, Queensland, Australia, was obtained from DAF Biological Collections [11]. The isolate was cultured on potato dextrose agar (Difco) and incubated at 23 to 25 °C for two months. Whole genomic DNA was extracted using the DNeasy Plant Mini kit (QIAGEN) according to the manufacturer’s protocol. Paired-end libraries were prepared according to Illumina Nextera^TM^ DNA Flex Library Prep Reference Guide using a Nextera^TM^ DNA Flex Library Prep Kit and Nextera^TM^ DNA CD Indexes. WGS sequencing was performed on Illumina MiSeq platform (600-cycles) at the molecular laboratories of the Centre for Crop Health, USQ. Assembly was performed on the Galaxy-Melbourne/GVL 4.0.0 webserver [83]. Raw reads were quality checked using FastQC (v0.11.5) [84] and trimmed using Trimmomatic (v0.36) [85] with the following parameters: TruSeq3 adapter sequences were removed using default settings, reads were cropped to remove 20 bases from the leading end and 65 bases from the trailing end of each read, minimum quality of leading and trailing bases was set to 30, a sliding window of four bases was used to retain those with an average quality of 30 and the minimum length read retained was 31 bases. *De novo* assembly was performed in two steps, first using Velvet (v1.2.10) [86] and VelvetOptimiser (v2.2.5) [87] with input k-mer size range of 81-101 (step size of 2). Secondly, SPAdes (v3.11.1) [88] was run on trimmed reads with the following parameters: read error correction, careful correction, automatic k-mer values, automatic coverage cutoff and Velvet contigs (>500 bp in length), from the previous step, included as trusted contigs. Contigs >500 bp in length were retained. Reads were mapped back to the assembly using Bowtie2 (v2.2.4) [89] and Picard toolkit (v2.7.1) [90] and visualised using IGV (v2.3.92) [91]. The genome assembly was checked for completeness with BUSCO (v2.0) [92] using the Ascomycota orthoDB (v9) dataset [93]. The extent and location of AT-rich regions was determined using OcculterCut (v1.1) [94] with default parameters and mitochondrial contigs.

The prediction of genes and transposable elements (TE) was performed on the GenSAS (v6.0) web platform [95], using GeneMarkES (v4.33) [96] for gene prediction and RepeatMasker (v4.0.7) [97], using the NCBI search engine and slow speed sensitivity, for the prediction of TE’s. Predicted gene models containing short exons, missing a start or stop codon or which overlapped a TE region were removed from the predicted proteome. The genome was searched for Short Simple Repeats (SSR) using the Microsatellite Identification tool (MISA) [98], with the SSR motif minimum length parameters being 10 for mono, 6 for di, and 5 for tri, tetra, penta and hexa motifs.

Annotation was performed using BLASTP (v2.7.1+) [99] to query the *E. fawcettii* predicted proteome against the Swiss-Prot Ascomycota database (release 2018_08) [100] with an e-value of 1e-06 and word size of 3. BLAST results were loaded into Blast2GO Basic (v5.2.1) [101], with InterProScan, mapping and annotation steps being performed with default parameters, except HSP-hit coverage cutoff was set to 50% to increase stringency during annotation. Further annotation was achieved using HmmScan in HMMER (v3.2.1) [102] to query the predicted proteome against the Protein Family Database (Pfam) (release 32) [103]. GC% content of the coding DNA sequence (CDS) of each gene was determined using nucBed from Bedtools (v2.27.1) [104]. Predicted proteins were searched for polyamino acid (polyAA) repeats of at least five consecutive amino acid residues using the FIMO motif search tool [105] within the Meme suite (v5.0.2) [106]. The Whole Genome Shotgun project was deposited at DDBJ/ENA/GenBank under the accession SDJM00000000. The version described in this paper is version SDJM01000000. Raw reads were deposited under the SRA accession PRJNA496356.

### Phylogenetic Analysis

ITS and partial TEF1α sequences of 12 *E. fawcettii* pathotypes, 11 closely related *Elsinoë* species and *Myriangium hispanicum* were obtained from GenBank (accessions provided in S1) for phylogenetic analysis with *E. fawcettii* (BRIP 53147a). Sequences for each locus were aligned using MUSCLE [107] with a gap open penalty of -400, concatenated and used to perform maximum likelihood analysis in MEGA7 [108] based on the General Time Reversible model [109] with partial deletion of 90% and 1000 bootstrap replicates. The initial tree for the maximum likelihood analysis was automatically selected using Neighbor-Join and BioNJ on the matrix of pairwise distances estimated using the Maximum Composite Likelihood method. A discrete Gamma distribution utilising 4 categories (+G, parameter = 0.4095) was used and the rate variation model allowed some sites to be invariable (+*I*, 26.6862% sites). The character matrix and tree were combined and converted to nexus format using Mesquite (v3.6) [110] prior to TreeBASE submission (TreeBASE reviewer access: http://purl.org/phylo/treebase/phylows/study/TB2:S25460?x-access-code=f3c2b3e55c147986b2a24b44407d9e48&format=html). *E. fawcettii* (BRIP 53147a) ITS and partial TEF1α sequences (accessions MN784182 and MN787508) were submitted to GenBank.

### Sequence Information

Genome assemblies and predicted proteomes included in the comparative analysis were obtained from GenBank. These included *U. maydis* (accession GCF_000328475.2, no. of scaffolds = 27) [111], *L. maculans* (accession GCF_000230375.1, no. of scaffolds = 76) [112], *M. oryzae* (accession GCF_000002495.2, no. of scaffolds = 53) [113], *R. commune* (accession GCA_900074885.1, no. of scaffolds = 164) [114], *V. dahliae* (accession GCF_000150675.1, no. of scaffolds = 55) [115], *B. cinerea* (accession GCF_000143535.2, no. of scaffolds = 18) [116], *Parastagonospora nodorum* (accession GCF_000146915.1, no. of scaffolds = 108) [117], *Pyrenophora tritici-repentis* (accession GCA_003231415.1, no. of scaffolds = 3964) [118], *S. sclerotiorum* (accession GCF_000146945.2, no. of scaffolds = 37) [119] and *Z. tritici* (accession GCA_900184115.1, no. of scaffolds = 20) [120]. Sequences of experimentally verified effector proteins were obtained from EffectorP 2.0 [121]. TE’s were identified in each assembly, as described above for *E. fawcettii*, and predicted genes which overlapped them were similarly removed from predicted proteomes.

### Prediction of secretome and effectors

Secretome and effector prediction was performed on the predicted proteomes of *E. fawcettii* and 10 fungal species known to contain effector proteins. Secretome prediction for each species began with a set of proteins predicted as secreted by either SignalP (v4.1) [122], Phobius [123] or ProtComp-AN (v6) [124]. This set was run through both the TMHMM Server (v2.0) [125] and PredGPI [126] to predict proteins with transmembrane helices and GPI-anchors, respectively. Those proteins with >1 helix or with 1 helix beyond the first 60 amino acids were removed, as were those with “highly probable” or “probable” GPI anchors. Remaining proteins formed the predicted secretome and were subjected to candidate effector prediction using EffectorP (v2.0) [121].

### Genomic, proteomic and known effector analyses

Sequences of 42 experimentally verified effector proteins, which showed >98% similarity to proteins from the 10 species included in this study, and which appeared in both the predicted secretome and candidate effector list for the respective species, were utilised in the known effector analysis. The following analyses were performed on the proteome/genome of each species. Results relating to the 42 known effectors were compared to results of all proteins from each species. Length of the intergenic flanking region (IFR) was determined as the number of bases between the CDS of two adjacent genes. Median IFR values were determined in R (v3.5.1) [127]. Genes were labelled as gene-dense if the IFR on each side was less than the median IFR length for that particular species, genes on a contig edge were not included among gene-dense labelled genes. Genes with IFR greater than the median on both sides were labelled as gene-sparse. SM clusters were predicted by passing genome assemblies and annotation files through antiSMASH fungal (v4.2.0) [128] using the Known Cluster Blast setting. Core, accessory and unique genes for each species were determined by mapping proteins into ortholog groups using the orthoMCL algorithm [129] followed by ProteinOrtho (v5.16b) [130] on remaining unclassified genes. Core genes were those shared by all comparative species, accessory genes were shared by at least two species, but not all, and unique genes were found in only one species. GC% content of the CDS of each gene was determined as described above, Q_1_ and Q_3_ values were determined for each species using R [127]. HmmScan [102] of all protein sequences against the Pfam database [103] was performed as described above. Genomic AT-rich region identification was performed using OcculterCut (v1.1) [94] as described above. For genomes with identified AT-rich regions, the distance between genes and their closest AT-rich region edge was determined using Bedtools closestBed [104], as was the distance between genes and the closest TE.

### Prioritisation of candidate effectors

CE’s of each species were prioritised using an optimised scoring system based on the analysis of known effectors in 10 fungal species. All were scored out of at least five points, corresponding to one point allocated for each of the following conditions: (I) not labelled as gene-dense; (II) no involvement in predicted SM clusters; (III) labelled as either unique to the species or allocated to the same orthoMCL group as a known effector; (IV) GC% of CDS was either below the Q_1_ value or above the Q_3_ value of the respective species; and (V) within 10 genes upstream or downstream was at least one gene coding for a protein with a top Pfam ID hit from the following list: p450, Mito carr, FAD binding 3, FAD binding 4, Ras, DUF3328, BTB, Peptidase M28, AA permease or AA permease 2. For species with genomes which had >2% TE coverage or >25% AT-rich region coverage, CE’s were scored out of six points. Those genomes which had both >2% TE and >25% AT-rich region coverage, CE’s were scored out of seven points. Hence, all candidate effectors were scored out of *n* (five, six or seven) points, those CE’s which obtained a score of *n* or *n*-1 points were labelled as prioritised CE’s.

### Prediction of other virulence genes

SM clusters were predicted using antiSMASH fungal (v4.2.0) [128] as described above. CAZymes were predicted by passing the predicted proteomes through the dbCAN2 meta server [131] and selecting three tools including HMMER scan against the dbCAN HMM database [132], Diamond [133] search against the Carbohydrate-Active enZYmes (CAZy) database [134] and Hotpep query against the Peptide Pattern Recognition library [135]. Predicted CAZymes were taken as those with positive results for at least two out of the three tools. Potential pathogenesis-related proteins were identified by querying the predicted proteomes against the Pathogen Host Interactions Database (PHI-base) (v4.6, release Oct 2018) [136] using BlastP (v2.7.1) [99] analyses with an e-value of 1e-06 and a query coverage hsp of 70%, those results with >40% similarity were retained. Prioritised candidate CWDE’s were shortlisted from the predicted CAZymes to those which were predicted as secreted and obtained hits to plant associated fungal pathogenicity-related genes in PHI-base which showed evidence of reduced virulence in knockout or mutant experiments.

## Results and Discussion

### Genome assembly and features

The genome assembly of *E. fawcettii* (BRIP 53147a), deposited at DDBJ/ENA/GenBank (accession SDJM00000000), was sequenced using paired-end Illumina WGS sequencing technology. Assembly of reads produced a draft genome 26.01 Mb in size with a coverage of 193x (Table 1) and consisted of 286 contigs greater than 500 bp in length, with an N50 of 662,293 bp, a mean contig length of 90,948 bp and an overall GC content of 52.3%. Running the assembly against the Ascomycota orthoDB (v9) [93] showed 97.6% of complete single copy genes were found in the *E. fawcettii* assembly, indicating a high degree of coding DNA sequence completeness. The genome of *E. fawcettii* is comparable in size to other fungal genomes including *Eurotium rubrum* (26.21 Mb) [137], *Xylona heveae* (24.34 Mb) [138] and *Acidomyces richmondensis* (29.3 Mb) [139], however it is smaller than the average Ascomycota genome size of 36.91 Mb [140]. When analysed against the 10 fungal species included in this comparative analysis (*B. cinerea*, *L. maculans*, *M. oryzae*, *Parastagonospora nodorum*, *Pyrenophora tritici-repentis*, *R. commune*, *V. dahliae, S. sclerotiorum*, *U. maydis* and *Z. tritici*), the *E. fawcettii* assembly is the second smallest, after *U. maydis* at 19.6 Kb. TE identification, by analysis against Repbase (release 18.02) [141], showed a coverage of only 0.37%, indicating a low proportion of the *E. fawcettii* genome is represented by currently known TE’s, this is a likely contributor to its comparatively small genome size. This low TE coverage may also be the result of a fragmented genome [142]. It is possible, should long read sequencing of this isolate be completed in the future, TE coverage may appear higher.

**Table 1.** Features of *Elsinoë fawcettii* (BRIP 53147a) genome assembly

| General Features |  |
| --- | --- |
| Assembly length (bp) | 26,011,141 |
| Coverage | 193x |
| Number of contigs | 286 |
| Mean GC content (%) | 52.3 |
| N50 (bp) | 694,004 |
| Mean contig length (bp) | 90,948 |
| Minimum contig length (bp) | 501 |
| Maximum contig length (bp) | 2,345,732 |
| Coverage of interspersed repeats (bp) | 95,654 (0.37%) |
| Coverage of short simple repeats (bp) | 6868 (0.026%) |
| Number of predicted gene models | 10,080 |
| Number of contigs containing predicted genes | 141 |
| Mean gene length (bp) | 1,573 |
| Mean number of exons per gene | 2.35 |
| Number of genes containing a polyAA repeat | 1,073 |
| Mean GC content of CDS (%) | 54.7 |

The *E. fawcettii* genome has less predicted gene models than the average Ascomycota genome of 11129.45 [140]. Gene prediction produced 10,080 gene models, 5,636 (55.91%) of which were annotated, while 4,444 (44.09%) were labelled as coding for hypothetical proteins. The average gene length was 1,573 bp with an average of 2.35 exons per gene, there were 3,280 single exon genes. The mean GC content of CDS was 54.7%, which was 2.4% higher than the overall GC content and showed a wide variation in range, with the lowest scoring gene at 44.29% GC and the highest being 71.53%, thus exposing a spectrum on which genes may be differentiated. Hmmscan [102] analysis of the predicted proteome against the Pfam database [103] revealed a high proportion (70.1% = 7,069) of genes with at least one hit to a Pfam model. The same analysis performed on the proteomes of the 10 fungal species included in the comparative analysis gave results ranging from 48.6% for *S. sclerotiorum*, with the lowest proportion of Pfam hits, to 74.9% for *U. maydis* with the highest, and a mean of 62.1% over the 11 species (S2).

Analysis of orthologous genes among *E. fawcettii* and the 10 comparative species indicated 3,077 (30.5%) of the predicted genes of *E. fawcettii* were core genes, finding hits through OrthoMCL or ProteinOrtho in all 11 species (S2). There were 4,874 (48.4%) *E. fawcettii* genes found in at least one other species but not all and were therefore considered accessory genes. Lastly, the remaining 2,129 (21.1%) were found in only the *E. fawcettii* proteome, 140 of these, however, obtained a hit to an orthoMCL group and were therefore set aside and not considered as unique proteins in subsequent analyses, leaving 1,989 (19.7%) genes presumed to be *Elsinoë*-specific and therefore potentially involved in either *Elsinoë*- or *E. fawcettii*-specific pathogenesis pathways. A comparative analysis among the core, accessory and unique genes of the 11 species (S2) (Figure 1) indicated that *U. maydis* was set apart from the other species by showing the lowest proportion of accessory genes, this was expected as *U. maydis* was the only biotroph and Basidiomycete among the group. *E. fawcettii* showed a below average percentage of unique genes which may be expected due its smaller than average sized genome and proteome. A lower number of unique genes may place a limitation on the ability of *E. fawcettii* to infect a larger range of host plants.

**Figure 1.**
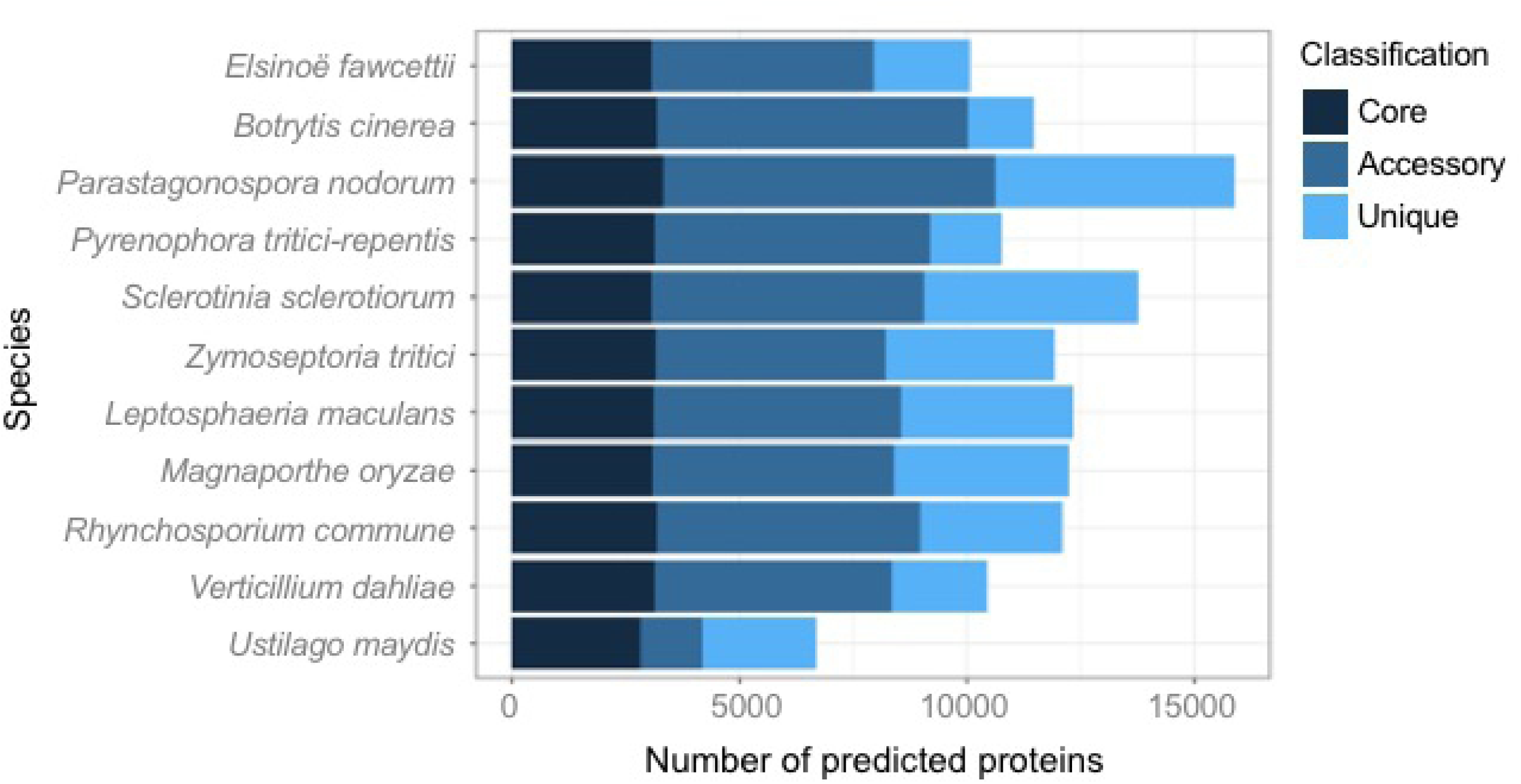
Comparison of gene classifications among the proteomes of 11 fungal pathogens. Genes were categorised using orthoMCL group IDs, or proteinortho if no group was assigned. Genes were considered; (I) core if they were shared by all 11 species; (II) accessory if they were shared by at least two species, but not all; (III) unique if they were found in only one of the 11 species.

While the overall GC content of *E. fawcettii* was 52.3%, when taking AT-rich regions into consideration, the average GC content of 98.97% of the genome was 52.8%, while the AT-rich regions had an average GC content of 33.8%. AT-rich regions are sections of DNA that are scattered throughout the genome and have a significantly higher AT content compared to adjacent GC equilibrated blocks [94]. The presence of AT-rich regions in genomes varies widely, for example *S. sclerotiorum* does not show evidence of AT-rich regions [143], while 36% of the *L. maculans* genome is covered by AT-rich regions which have an average GC content of 33.9% [47]. AT-rich regions are thought to develop in, and nearby to, regions containing TE repeats, through Repeat-Induced Point mutation (RIP), a mechanism used to inhibit the destructive actions of TE’s against an organism’s genome. Through a fungal genome defence mechanism causing cytosine to thymine polymorphisms, a TE repeat sequence is inhibited from further movement and potential destruction of necessary genes. This same type of polymorphism can also occur in genes nearby to TE regions [144–147], potentially providing numerous genomic locations with increased plasticity scattered throughout the genome. While RIP occurs during the sexual phase it has also been observed in asexual fungi and is thought to indicate a species reproductive history or potential [148]. AT-rich regions are present within the *E. fawcettii* genome, however the extent of their coverage in the present assembly is low, 59 regions with an average GC content of 33.8% cover only 1.03% of the genome. Sixteen regions are found overlapping TE’s, while four are found within 2 Kb of a TE region, meaning 33.9% of the AT-rich regions potentially represent RIP-affected regions. The remaining 66.1%, found either >2Kb away or on a contig that does not contain a predicted TE region, are potentially RIP-affected regions where the TE is no longer recognisable. The AT-rich regions of *E. fawcettii* are not scattered evenly throughout the genome, instead 29/59 (49.2%) are situated at the edge of a contig and 15/59 (25.4%) cover the entire length of a contig, specifically contigs not containing genes. Two further AT-rich regions were located between the edge of a contig and the beginning of the first gene and so were grouped with those located at the edge of a contig. The remaining 13 regions (22.0%) were situated within a contig with genes residing on both sides. Hence, the majority either made up the edge of a contig which contained genes or filled entire contigs which did not contain genes, meaning it is likely that the sequence of many *E. fawcettii* AT-rich regions contain sections of such low complexity that contig breaks result, a hypothesis which could be tested in the future using long read sequencing technology. Eight predicted genes at least partially overlap these regions and 57 are located within 2 Kb, a finding which has potential significance as AT-rich regions have been known to harbour effector genes in fungal pathogens [149, 150]. There was a large range of diversity of AT-rich region coverage among the fungal pathogens analysed in the current study; *S. sclerotiorum*, *Pyrenophora tritici-repentis*, *M. oryzae* and *U. maydis* showed no AT-rich regions; *V. dahliae* (1.5%), *B. cinerea* (4.9%), *Parastagonospora nodorum* (6.6%) and *Z. tritici* (17.3%) showed lower degrees of AT-rich coverage; while *R. commune* (29.5%) and *L. maculans* (37%) showed the greatest extent. These levels of AT-rich coverage did not appear to corelate with pathogen classification as necrotrophic, hemibiotrophic or biotrophic, nor as host-specific or broad-host range pathogens. The genomic location of AT-rich regions was, however, further included in the known effectors and candidate effectors analyses. Identification and analysis of SSR’s in the *E. fawcettii* genome located 400 regions covering 6,868 bp (0.026%), 164 (41%) of which were contained within a predicted gene.

Furthermore, polyAA repeats, of at least five identical and adjacent residues, were identified within 1,073 predicted protein sequences. The presence of repetitive sequences has been noted in fungal effectors [33, 45, 151] and implicated in the function and evolution of pathogenicity-related genes of other plant-associated microorganisms [152]. Analysis of the 1,105 proteins which obtained either an SSR or polyAA hit indicated 237 (21.45%) were categorised as *E. fawcettii*-specific and did not obtain a Pfam hit, highlighting potentially novel genus- or species-specific genes involved in host pathogenesis.

Phylogenetic analysis of partial ITS and TEF1-α regions of *E. fawcettii* (BRIP 53147a) in comparison with other *E. fawcettii* isolates and closely related *Elsinoë* species (Figure 2) indicates *E. fawcettii* (BRIP 53147a) closely aligns with the *E. fawcettii* clade. Substitutions appearing in the Jingeul pathotype isolates are not seen in isolate BRIP 53147a. One G to A substitution in the TEF1-α region sets isolate BRIP 53147a apart from the other *E. fawcettii* isolates (S3), a base which is at the 3^rd^ position of a Glu codon and hence does not result in a translational difference. This substitution in the BRIP 52147a isolate appeared with a high degree of confidence, 100% of sequence reads aligned back to the assembly and a coverage of 241x, at this point, agreed with the substitution. While it is thought that isolate BRIP 53147a belongs to either the Lemon or Tyron’s pathotype, it is yet to be determined which or if it constitutes a new pathotype of its own. Aside from the one base substitution in the TEF1-α region, there would be some expected differences throughout the genomes of the *E. fawcettii* BRIP 53147a isolate and the other *E. fawcettii* isolates due to differences in collection details, such as geographical location, year and host specificity. Specifically, isolate BRIP 53147a was collected in Montville, Queensland in 2009, while the other Australian isolates, DAR 70187 and DAR 70024, belonging to the Lemon and Tyron’s pathotypes, were collected 15 years earlier in Somersby and Narara in NSW, respectively [7], both a distance of almost 1000 km away. Several isolates from Figure 2 have been tested for host pathogenicity leading to the designation of specific pathotypes [3], as opposed to relying on only sequence data and thus illustrating the importance of experimental validation prior to pathotype or species classification. For example, Jin-1 and Jin-6 are classified as the Jingeul pathotype, SM3-1 as FBHR, S38162 as FNHR, CC-132 as SRGC, DAR 70187 and CC-3 as the Lemon pathotype, and DAR 70024 as Tyron’s pathotype [3]. Host specificity experimentation for the *E. fawcettii* BRIP 53147a isolate is a suggested future step, as is the whole genome sequencing and analysis of further *E. fawcettii* isolates for comparison. The comprehensive host pathogenicity testing of 61 *E. fawcettii* isolates and their subsequent classification into six pathotypes [3] coupled with genomic sequencing data analysis would provide a wealth of knowledge of potential host-specific pathogenicity-related genes and mutations.

**Figure 2.**
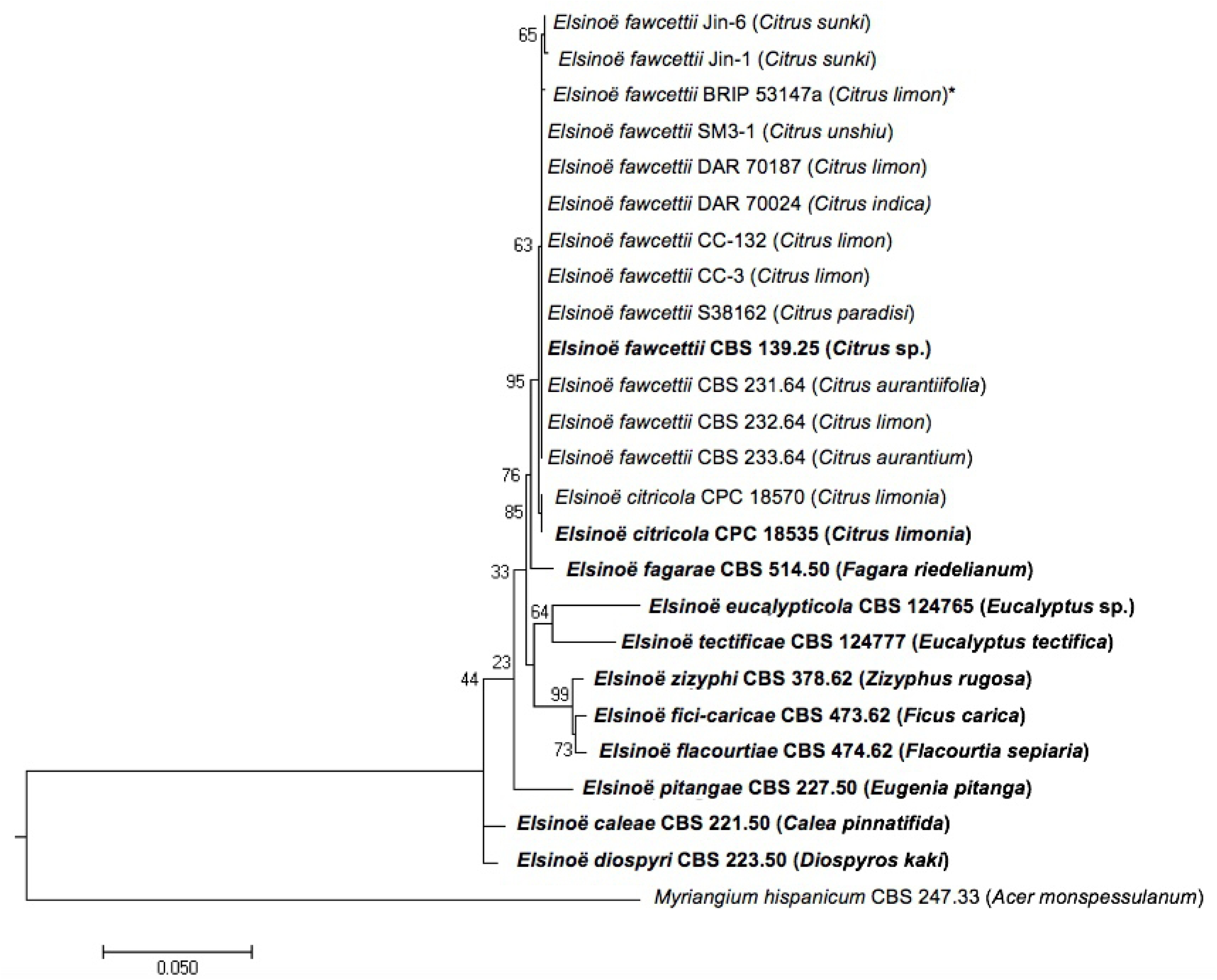
Maximum likelihood phylogenetic tree of *E. fawcettii* isolates and closely related species. The phylogenetic tree was inferred from a concatenated dataset including ITS and partial TEF1-α regions. *Myriangium hispanicum* was used as the outgroup. The branch length indicates the number of nucleotide substitutions per site, bootstrap values are shown at nodes, host in parentheses, new isolate described in the current study denoted with asterisk (*) and type strains are in bold.

### Prediction of secretome and effectors

A total of 1,280 genes (12.7% of the proteome) were predicted to code for secreted proteins (SP) in the *E. fawcettii* genome (Table 2). Using the discovery pipeline outlined in Figure 3, classically secreted proteins with a detectable signal peptide were predicted by either SignalP and/or Phobius providing 1,449 proteins, while ProtComp identified a further 120 as potential non-classically secreted proteins. Of these 1,569 proteins 186 were removed as they were predicted to contain transmembrane helices, an indication that while targeted for secretion the protein likely functions while situated in the cell membrane. A further 103 were removed as they contained a predictable GPI anchor, also suggesting they associate with the cell membrane to perform their function, leaving a total of 1,280 proteins identified as likely SP’s. To enable comparison of the species’ predicted secretomes and CE’s, the same prediction pipeline (Figure 3) was used on the proteomes of 10 further fungal species included in the analysis (Table 2), essentially utilising genomes which contain known protein effectors for comparison. The proportion of predicted SP’s in the *E. fawcettii* proteome was similar to that of other necrotrophic fungal pathogens, which ranged from *B. cinerea* at 11.3% to *Parastagonospora nodorum* at 13.9%. It was, however, lower in comparison to the hemibiotrophs; *R. commune* showed a low of 12.5% SP’s while *M. oryzae* was the highest scoring at 18.5%, demonstrating a small increase in proportion of SP’s for the hemibiotrophs compared to the necrotrophs. This potentially provides them with a larger array of secreted proteins compared to biotrophs and necrotrophs, to first support a biotrophic, and secondly a necrotrophic, host interaction.

**Figure 3.**
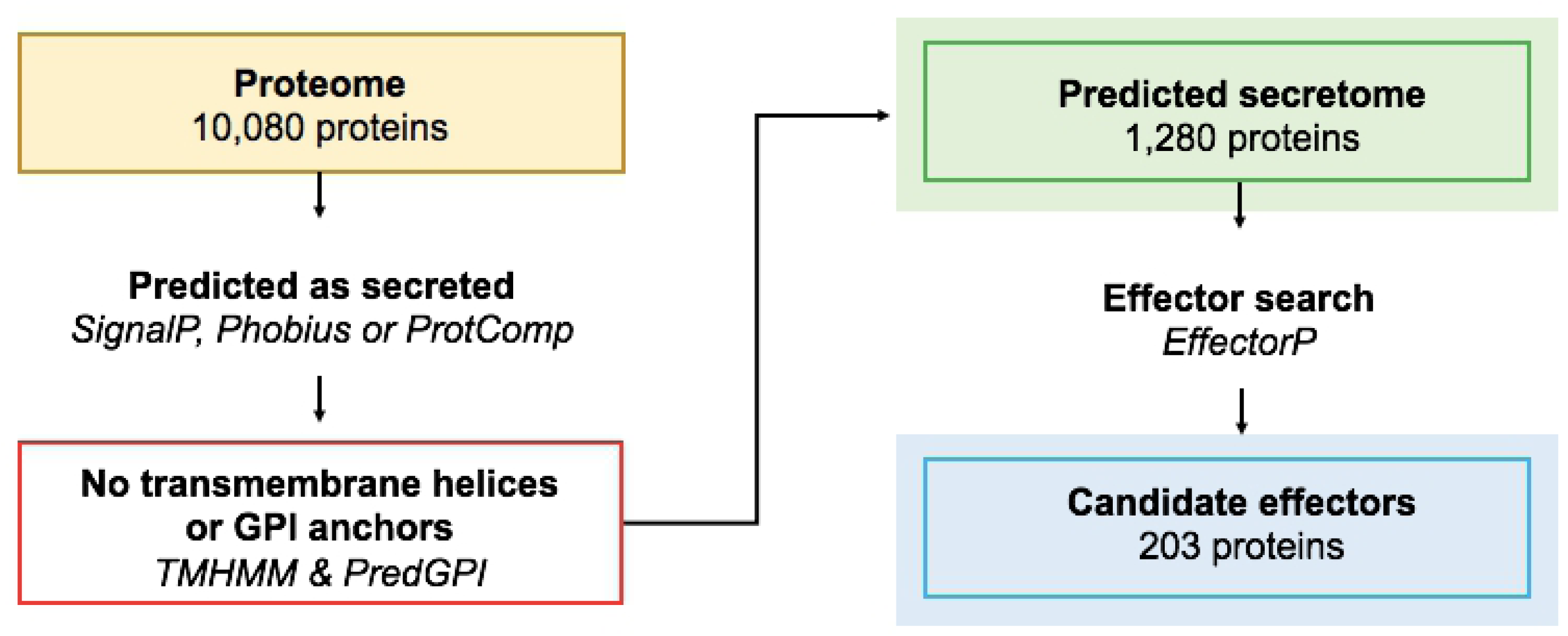
Pipeline for the discovery of the predicted secretome and candidate effectors. The secretome search started with the predicted proteins of a species, proteins were predicted as secreted using at least one of three tools, proteins with predicted transmembrane helices or GPI-anchors were removed. Candidate effectors were predicted using EffectorP. The number of proteins shown for the predicted proteome, secretome and effectome refers to the *Elsinoë fawcettii* BRIP 53147a genome.

**Table 2.**
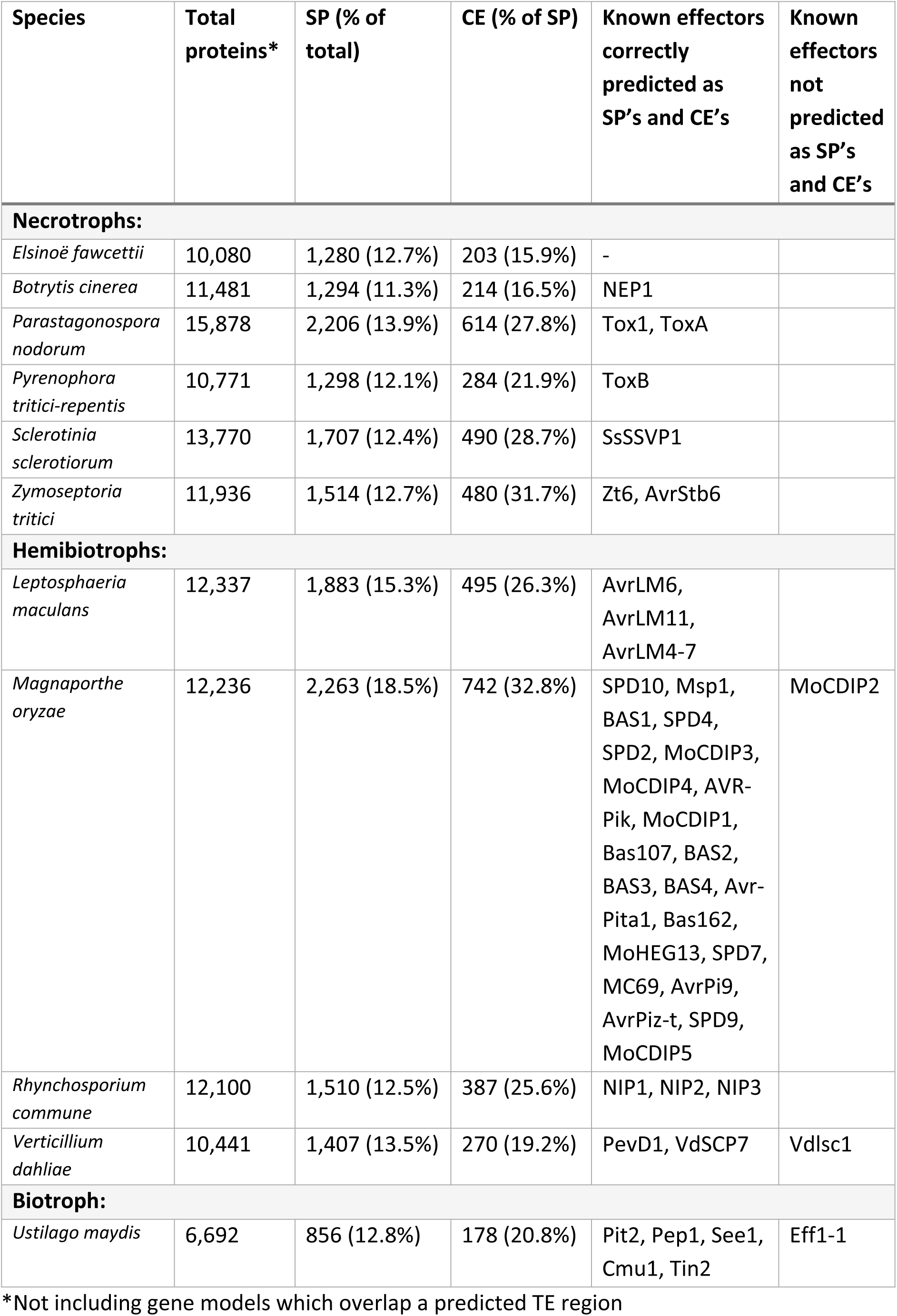
Predicted secreted proteins, candidate effectors and known effectors

| Species | Total proteins* | SP (% of total) | CE (% of SP) | Known effectors correctly predicted as SP's and CE's | Known effectors not predicted as SP's and CE's |
| --- | --- | --- | --- | --- | --- |
| **Necrotrophs:** |  |  |  |  |  |
| *Elsinoë fawcettii* | 10,080 | 1,280 (12.7%) | 203 (15.9%) | - |  |
| *Botrytis cinerea* | 11,481 | 1,294 (11.3%) | 214 (16.5%) | NEP1 |  |
| *Parastagonospora nodorum* | 15,878 | 2,206 (13.9%) | 614 (27.8%) | Tox1, ToxA |  |
| *Pyrenophora tritici-repentis* | 10,771 | 1,298 (12.1%) | 284 (21.9%) | ToxB |  |
| *Sclerotinia sclerotiorum* | 13,770 | 1,707 (12.4%) | 490 (28.7%) | SsSSVP1 |  |
| *Zymoseptoria tritici* | 11,936 | 1,514 (12.7%) | 480 (31.7%) | Zt6, AvrStb6 |  |
| **Hemibiotrophs:** |  |  |  |  |  |
| *Leptosphaeria maculans* | 12,337 | 1,883 (15.3%) | 495 (26.3%) | AvrLM6, AvrLM11, AvrLM4-7 |  |
| *Magnaporthe oryzae* | 12,236 | 2,263 (18.5%) | 742 (32.8%) | SPD10, Msp1, BAS1, SPD4, SPD2, MoCDIP3, MoCDIP4, AVR-Pik, MoCDIP1, Bas107, BAS2, BAS3, BAS4, Avr-Pita1, Bas162, MoHEG13, SPD7, MC69, AvrPi9, AvrPiz-t, SPD9, MoCDIP5 | MoCDIP2 |
| *Rhynchosporium commune* | 12,100 | 1,510 (12.5%) | 387 (25.6%) | NIP1, NIP2, NIP3 |  |
| *Verticillium dahliae* | 10,441 | 1,407 (13.5%) | 270 (19.2%) | PevD1, VdSCP7 | Vdlsc1 |
| **Biotroph:** |  |  |  |  |  |
| *Ustilago maydis* | 6,692 | 856 (12.8%) | 178 (20.8%) | Pit2, Pep1, See1, Cmu1, Tin2 | Eff1-1 |

507 \*Not including gene models which overlap a predicted TE region

Known effectors were frequently identified by the CE pipeline (Figure 3), with 43/45 (95.6%) correctly predicted as being secreted and 42/45 (93.3%) also predicted as effectors (Table 2), highlighting the effectiveness of the pipeline among these fungal species. Those known effectors which were tested but not identified as SP’s included Vdlsc1 (*V. dahliae*) and MoCDIP2 (*M. oryzae*). Vdlsc1 lacks an N-terminal signal peptide and is unconventionally secreted [153], however it was not identified as a non-classically secreted protein. MoCDIP2 was removed as it obtained a GPI-anchor hit. Additionally, Eff1-1 (*U. maydis*) was predicted as secreted but not as a candidate effector, Eff1-1, along with MoCDIP2, are both known false negatives of EffectorP 2.0 [121].

The total number of CE’s identified for *E. fawcettii* was 203, meaning only 15.9% of SP’s gained CE classification, this was the lowest proportion out of all 11 species analysed (Table 2). This may be explained by the potential favouring of EffectorP towards SP’s of species on which it was trained. To further investigate this potential, results of EffectorP for the 11 species were compared to the results of an alternate candidate effector search; SP’s with a protein length less than the species’ median and with no Pfam hit other than to that of a known effector (S4). While this second method resulted in the identification of a higher number of CE’s for each species, *E. fawcettii* still obtained the lowest proportion of CE’s out of predicted SP’s, indicating *E. fawcettii* may have a lighter dependence, compared to other fungal pathogens, on protein effectors. It also highlighted the advantage of using EffectorP to narrow down an extensive catalogue of SP’s, as opposed to identifying CE’s based on arbitrary features. However, the CE’s predicted by EffectorP still range in the hundreds (Table 2), it was therefore beneficial to further shortlist candidates for prioritisation. To achieve this, known effectors which were correctly predicted as both SP’s and as CE’s (Table 2) were retained for further analysis to generate an optimised prioritisation scoring system.

### Known effector analysis

A total of 42 known effectors from 10 fungal species were analysed for; (I) gene density; (II) GC content; (III) involvement in SM clusters; (IV) uniqueness; (V) Pfam hits of surrounding genes; (VI) distance to the closest TE; and (VII) distance to the closest AT-rich regions (Table 3). Results were compared to those of all predicted genes from each of the same 10 species (S5). Features observed at a higher rate among the known effector group compared with each species’ proteome were used to generate a prioritisation pathway using a point allocation system. (I) Genes were labelled as gene-dense if the IFR’s on both sides were less than the IFR median value for that specific species, allowing an analysis relative to each organism. The proportions of gene-dense genes ranged from 21.6% (*Pyrenophora tritici-repentis*) to 28.0% (*B. cinerea*) (S5), in contrast to 3/42 (7.1%) known effectors (Table 3).

**Table 3.**
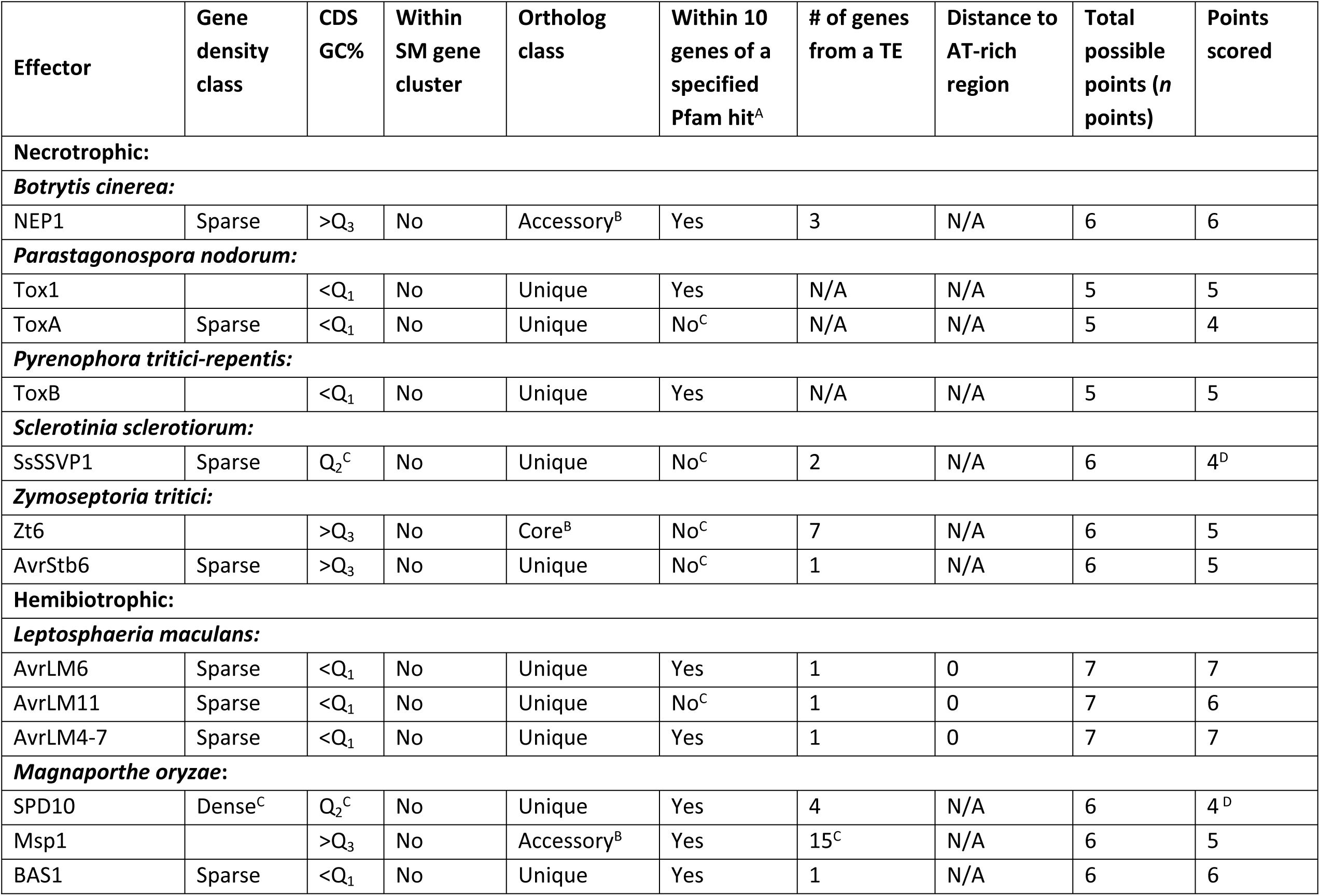
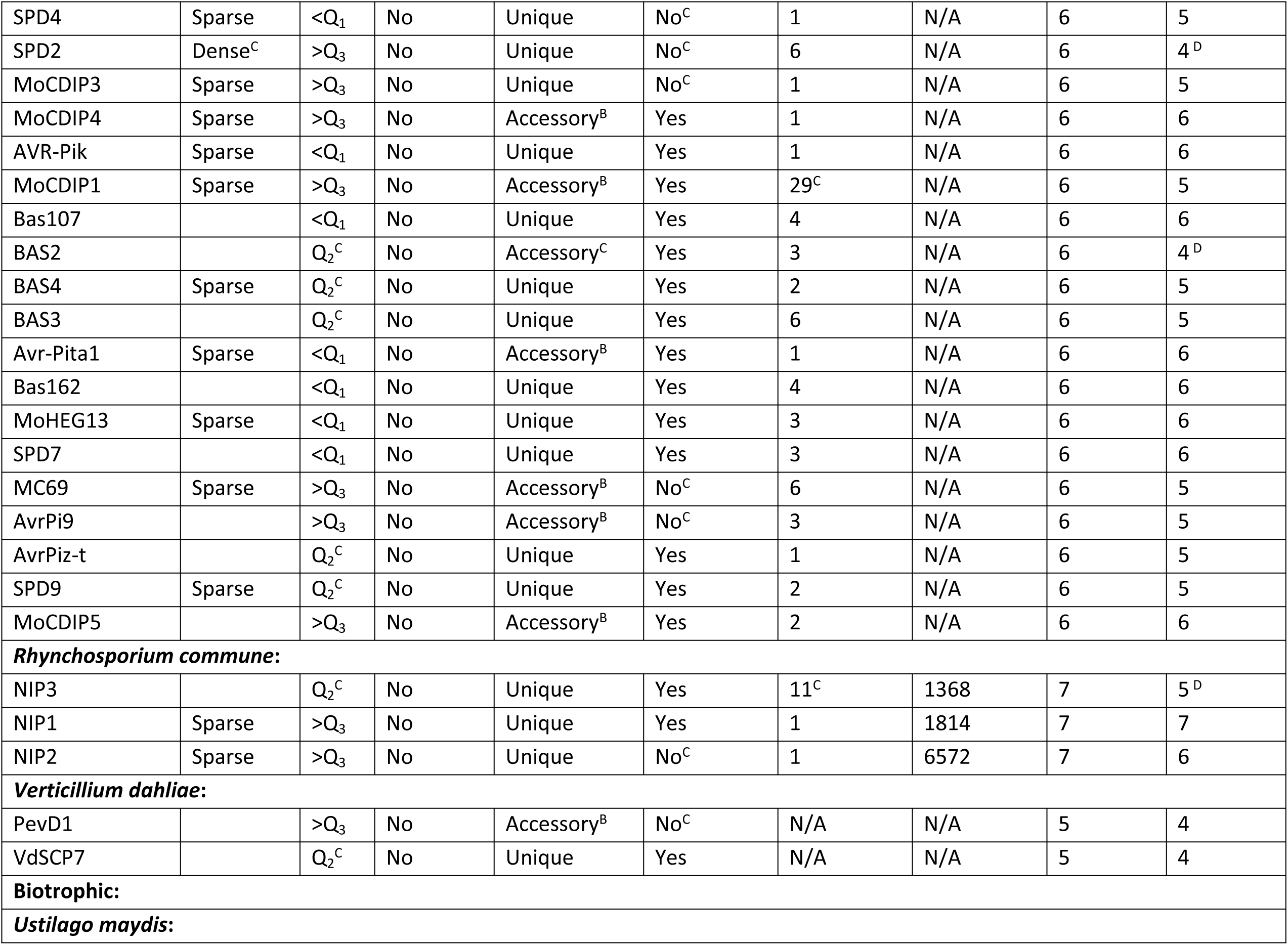
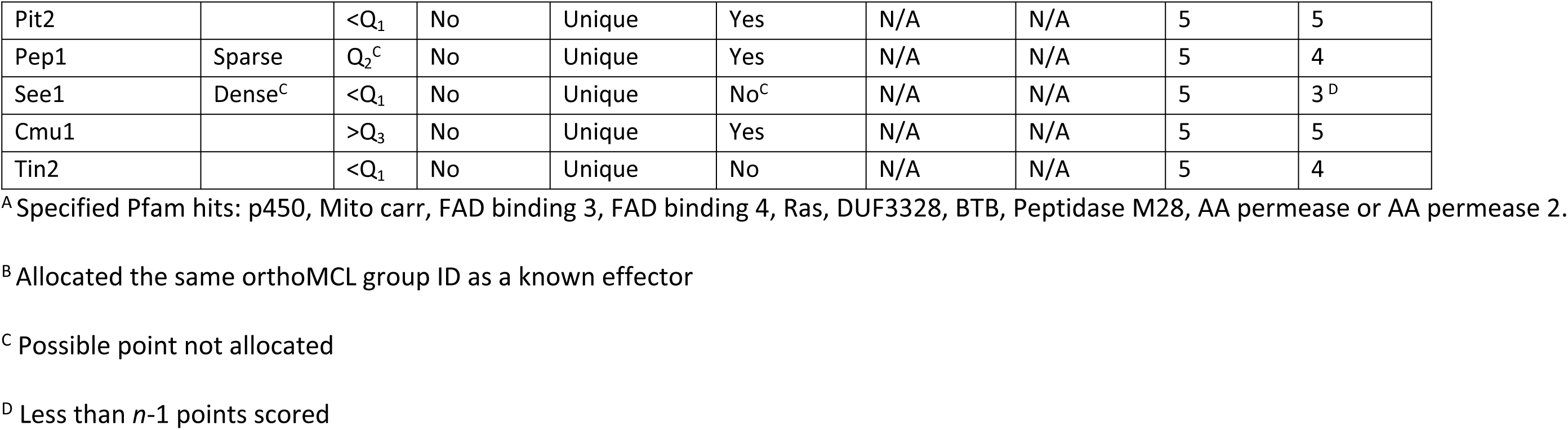
Features of known fungal effectors used to guide candidate effector prioritisation

| Effector | Gene density class | CDS GC% | Within SM gene cluster | Ortholog class | Within 10 genes of a specified Pfam hit<sup>A</sup> | # of genes from a TE | Distance to AT-rich region | Total possible points (*n* points) | Points scored |
| --- | --- | --- | --- | --- | --- | --- | --- | --- | --- |
| **Necrotrophic:** |  |  |  |  |  |  |  |  |  |
| ***Botrytis cinerea*:** |  |  |  |  |  |  |  |  |  |
| NEP1 | Sparse | >Q<sub>3</sub> | No | Accessory<sup>B</sup> | Yes | 3 | N/A | 6 | 6 |
| ***Parastagonospora nodorum*:** |  |  |  |  |  |  |  |  |  |
| Tox1 |  | <Q<sub>1</sub> | No | Unique | Yes | N/A | N/A | 5 | 5 |
| ToxA | Sparse | <Q<sub>1</sub> | No | Unique | No<sup>C</sup> | N/A | N/A | 5 | 4 |
| ***Pyrenophora tritici-repentis*:** |  |  |  |  |  |  |  |  |  |
| ToxB |  | <Q<sub>1</sub> | No | Unique | Yes | N/A | N/A | 5 | 5 |
| ***Sclerotinia sclerotiorum*:** |  |  |  |  |  |  |  |  |  |
| SsSSVP1 | Sparse | Q<sub>2</sub><sup>C</sup> | No | Unique | No<sup>C</sup> | 2 | N/A | 6 | 4<sup>D</sup> |
| ***Zymoseptoria tritici*:** |  |  |  |  |  |  |  |  |  |
| Zt6 |  | >Q<sub>3</sub> | No | Core<sup>B</sup> | No<sup>C</sup> | 7 | N/A | 6 | 5 |
| AvrStb6 | Sparse | >Q<sub>3</sub> | No | Unique | No<sup>C</sup> | 1 | N/A | 6 | 5 |
| **Hemibiotrophic:** |  |  |  |  |  |  |  |  |  |
| ***Leptosphaeria maculans*:** |  |  |  |  |  |  |  |  |  |
| AvrLM6 | Sparse | <Q<sub>1</sub> | No | Unique | Yes | 1 | 0 | 7 | 7 |
| AvrLM11 | Sparse | <Q<sub>1</sub> | No | Unique | No<sup>C</sup> | 1 | 0 | 7 | 6 |
| AvrLM4-7 | Sparse | <Q<sub>1</sub> | No | Unique | Yes | 1 | 0 | 7 | 7 |
| ***Magnaporthe oryzae*:** |  |  |  |  |  |  |  |  |  |
| SPD10 | Dense<sup>C</sup> | Q<sub>2</sub><sup>C</sup> | No | Unique | Yes | 4 | N/A | 6 | 4<sup>D</sup> |
| Msp1 |  | >Q<sub>3</sub> | No | Accessory<sup>B</sup> | Yes | 15<sup>C</sup> | N/A | 6 | 5 |
| BAS1 | Sparse | <Q<sub>1</sub> | No | Unique | Yes | 1 | N/A | 6 | 6 |
| SPD4 | Sparse | <Q<sub>1</sub> | No | Unique | No<sup>C</sup> | 1 | N/A | 6 | 5 |
| SPD2 | Dense<sup>C</sup> | >Q<sub>3</sub> | No | Unique | No<sup>C</sup> | 6 | N/A | 6 | 4<sup>D</sup> |
| MoCDIP3 | Sparse | >Q<sub>3</sub> | No | Unique | No<sup>C</sup> | 1 | N/A | 6 | 5 |
| MoCDIP4 | Sparse | >Q<sub>3</sub> | No | Accessory<sup>B</sup> | Yes | 1 | N/A | 6 | 6 |
| AVR-Pik | Sparse | <Q<sub>1</sub> | No | Unique | Yes | 1 | N/A | 6 | 6 |
| MoCDIP1 | Sparse | >Q<sub>3</sub> | No | Accessory<sup>B</sup> | Yes | 29<sup>C</sup> | N/A | 6 | 5 |
| Bas107 |  | <Q<sub>1</sub> | No | Unique | Yes | 4 | N/A | 6 | 6 |
| BAS2 |  | Q<sub>2</sub><sup>C</sup> | No | Accessory<sup>C</sup> | Yes | 3 | N/A | 6 | 4<sup>D</sup> |
| BAS4 | Sparse | Q<sub>2</sub><sup>C</sup> | No | Unique | Yes | 2 | N/A | 6 | 5 |
| BAS3 |  | Q<sub>2</sub><sup>C</sup> | No | Unique | Yes | 6 | N/A | 6 | 5 |
| Avr-Pita1 | Sparse | <Q<sub>1</sub> | No | Accessory<sup>B</sup> | Yes | 1 | N/A | 6 | 6 |
| Bas162 |  | <Q<sub>1</sub> | No | Unique | Yes | 4 | N/A | 6 | 6 |
| MoHEG13 | Sparse | <Q<sub>1</sub> | No | Unique | Yes | 3 | N/A | 6 | 6 |
| SPD7 |  | <Q<sub>1</sub> | No | Unique | Yes | 3 | N/A | 6 | 6 |
| MC69 | Sparse | >Q<sub>3</sub> | No | Accessory<sup>B</sup> | No<sup>C</sup> | 6 | N/A | 6 | 5 |
| AvrPi9 |  | >Q<sub>3</sub> | No | Accessory<sup>B</sup> | No<sup>C</sup> | 3 | N/A | 6 | 5 |
| AvrPiz-t |  | Q<sub>2</sub><sup>C</sup> | No | Unique | Yes | 1 | N/A | 6 | 5 |
| SPD9 | Sparse | Q<sub>2</sub><sup>C</sup> | No | Unique | Yes | 2 | N/A | 6 | 5 |
| MoCDIP5 |  | >Q<sub>3</sub> | No | Accessory<sup>B</sup> | Yes | 2 | N/A | 6 | 6 |
| ***Rhynchosporium commune:*** |  |  |  |  |  |  |  |  |  |
| NIP3 |  | Q<sub>2</sub><sup>C</sup> | No | Unique | Yes | 11<sup>C</sup> | 1368 | 7 | 5<sup>D</sup> |
| NIP1 | Sparse | >Q<sub>3</sub> | No | Unique | Yes | 1 | 1814 | 7 | 7 |
| NIP2 | Sparse | >Q<sub>3</sub> | No | Unique | No<sup>C</sup> | 1 | 6572 | 7 | 6 |
| ***Verticillium dahliae:*** |  |  |  |  |  |  |  |  |  |
| PevD1 |  | >Q<sub>3</sub> | No | Accessory<sup>B</sup> | No<sup>C</sup> | N/A | N/A | 5 | 4 |
| VdSCP7 |  | Q<sub>2</sub><sup>C</sup> | No | Unique | Yes | N/A | N/A | 5 | 4 |
| **Biotrophic:** |  |  |  |  |  |  |  |  |  |
| ***Ustilago maydis:*** |  |  |  |  |  |  |  |  |  |
| Pit2 |  | <Q<sub>1</sub> | No | Unique | Yes | N/A | N/A | 5 | 5 |
| Pep1 | Sparse | Q<sub>2</sub><sup>C</sup> | No | Unique | Yes | N/A | N/A | 5 | 4 |
| See1 | Dense<sup>C</sup> | <Q<sub>1</sub> | No | Unique | No<sup>C</sup> | N/A | N/A | 5 | 3<sup>D</sup> |
| Cmu1 |  | >Q<sub>3</sub> | No | Unique | Yes | N/A | N/A | 5 | 5 |
| Tin2 |  | <Q<sub>1</sub> | No | Unique | No | N/A | N/A | 5 | 4 |

<sup>A</sup> Specified Pfam hits: p450, Mito carr, FAD binding 3, FAD binding 4, Ras, DUF3328, BTB, Peptidase M28, AA permease or AA permease 2.
<sup>B</sup> Allocated the same orthoMCL group ID as a known effector
<sup>C</sup> Possible point not allocated
<sup>D</sup> Less than $n-1$ points scored

This provided grounds to allocate one point to each known effector which was not labelled as gene-dense. (II) GC content of the CDS of each gene was determined and median values calculated for each species, revealing the GC percentage of 32/42 (76.2%) known effectors fell either below the Q_1_ value or above the Q_3_ value for the respective species. When compared to an expected 50% in the upper and lower quartiles, this provided reason for the allocation of one point to known effectors should they fall in these two quartiles. (III) No overlap was observed between known effectors and the predicted SM clusters within each species, giving strong reason for the allocation of one point to known effector’s that were not included in SM clusters. (IV) Analysis of gene classification (core, accessory or unique) for each known effector highlighted that 41/42 (97.6%) were either unique to the species (31/42) or were assigned an orthoMCL group ID of a known effector (10/42). In contrast, the proportion of unique genes for each species was much lower, ranging from 11.9% (*B. cinerea*) to 33.7% (*S. sclerotiorum*), with an average of 25.4%. The proportion of genes allocated an orthoMCL of a known effector was similarly low at less than 0.3% for all species. Thus, a point was allocated to known effectors that were either unique to the species or obtained the same orthoMCL ID of a known effector. (V) Pfam hits of genes surrounding known effectors were also compared to the rates of Pfam hits from all 10 proteomes together. Analysis of the 10 genes upstream and downstream of a known effector revealed 10 Pfam hits which appeared at a rate at least double to that seen among the concatenated proteomes. For example, Pfam hits to cytochrome P450 accounted for 2.82% of all hits among the 10 genes up and downstream of a known effector, compared to only 1.26% of Pfam hits from the predicted proteins of all 10 species. Aside from cytochrome P450, further Pfam hits overrepresented among the genes surrounding known effectors included mitochondrial carrier protein, FAD binding domains 3 and 4, Ras family, domain of unknown function (DUF3328), BTB/POZ domain, peptidase family M28, and amino acid permease 1 and 2. At least one of these Pfam hits was found within 10 genes of 66.7% of the known effectors, which was higher when compared to all genes of each of the 10 species. Proportions ranged from only 28.7% (*Pyrenophora tritici-repentis*) up to 43.9% (*B. cinerea*), with an average of 34.1%, over the 10 species, of genes being within 10 genes of an overrepresented Pfam hit. A point was therefore allocated to known effectors which lay within 10 genes of a gene with one of the above mentioned Pfam hits. (VI) Those genomes with >2% TE coverage also showed a high proportion of known effectors in the close vicinity of TE’s. Specifically, 29/32 (90.6%) known effectors from *Z. tritici*, *S. sclerotiorum*, *B. cinerea*, *R. commune*, *L. maculans* and *M. oryzae* were within seven genes of a TE region, compared to an average of 47.8% of genes within seven genes of a TE for the same six species. This led to the allocation of one point for known effectors within seven genes of a TE for species with >2% TE coverage. (VII) Lastly, of the genomes analysed, only those consisting of >25% AT-rich regions, being *R. commune* and *L. maculans*, were found to have a noticeable association between the location of known effectors and AT-rich regions. The distance of all known effectors to the closest AT-rich region, of these two species, were found to be less than the Q_1_ value for each species. Hence, known effectors with these specifications, in species with >25% AT-rich region coverage, were allocated one point. It can be seen that depending on the degree of TE and AT-rich region coverage, each species known effectors may be scored out of five, six or seven points, henceforth referred to as “*n* points”. Over the 10 species with known effectors which were analysed, Table 3 illustrates a total of 36/42 (85.7%) known effectors obtained *n* or *n*-1 points, revealing a process which could be used to prioritise the many CE’s predicted for the *E. fawcettii* genome.

### Prioritisation of candidate effectors

While EffectorP correctly determined most known effectors, it also identified a large number of additional CE’s. While it is likely some of these candidates are unknown effectors being utilised by the pathogen to infect its host, it would be worthwhile to shortlist this group, to a list of the more likely candidates, prior to expensive and time-consuming experimental validation procedures. A points-based process was developed, based on the analysis of known effectors, to prioritise CE’s based on several features including: their distance to neighbouring genes, lack of involvement in predictable SM clusters, GC% of CDS, proximity to genes obtaining certain Pfam hits and potential uniqueness (Figure 4). For species with genome assemblies containing >2% TE coverage the number of genes a CE was from a TE was taken into consideration. Similarly, the distance between genes and AT-rich regions was acknowledged if AT-rich regions covered >25% of the species’ assembly. For each CE gene, one point was available for each of the above features, hence, as described for the known effector analysis, CE’s of each species were allocated a possible five, six or seven points (*n* points). *E. fawcettii*, *Parastagonospora nodorum*, *Pyrenophora tritici-repentis*, *V. dahlia* and *U. maydis* each had <2% TE coverage and <25% coverage of AT-rich regions, their CE’s were therefore scored out of five points. *Z. tritici*, *S. sclerotiorum*, *B. cinerea* and *M. oryzae* had >2% TE coverage but <25% coverage of AT-rich regions and so were scored out of six points. Only the assemblies of *R. commune* and *L. maculans* showed >2% TE’s and >25% AT-rich regions, and as such their CE’s were scored out of seven points. By using *n* or *n*-1 points as an acceptable score for CE prioritisation, revealed that CE’s of the 11 species could be reduced, by 51.1% - 83.6% (average 66.2%) (S6), with species that were scored out of more points achieving higher reductions.

**Figure 4.**
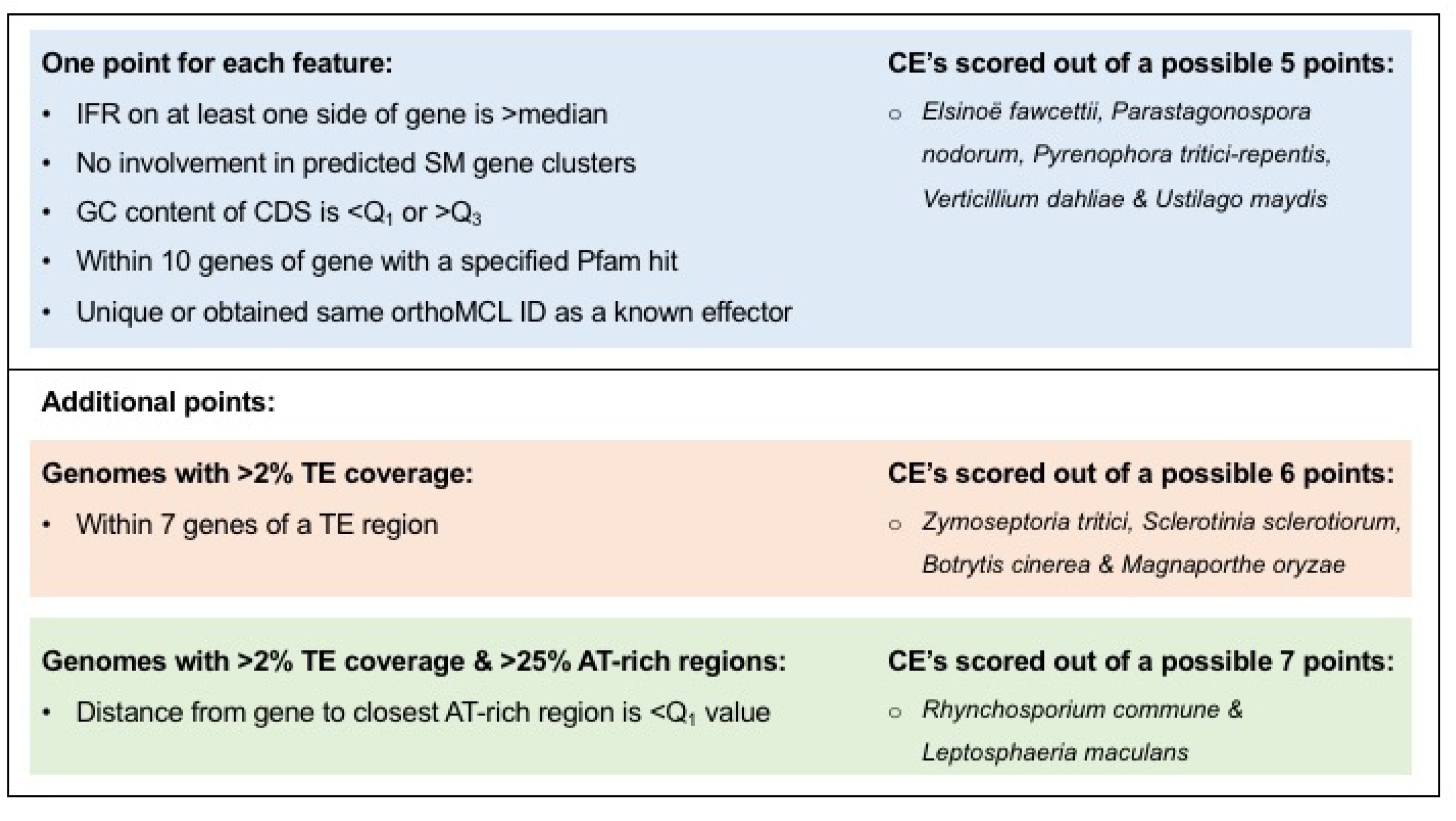
Candidate effector prioritisation features and points. The candidate effectors (CE’s) of all genomes analysed were scored using features shown in the blue box. Additional features were considered for CE’s from genomes with >2% TE coverage (red box) and >25% AT-rich region coverage (green box).

Applying the method outlined in Figure 4 to the CE’s of *E. fawcettii* led to the prioritisation of 77 CE’s, a reduction of 62%, for future experimental validation. This is a comparable reduction to that of the other necrotrophic pathogens (Figure 5, S6), for which six out of seven known effectors were retained within the shortlisted CE’s. Features of the 77 CE’s of *E. fawcettii* (S7) indicated many were small in size, had a high GC content, had a high proportion of cysteine residues and were more likely to be classified as gene-sparse. The median protein length was 181 aa, compared to 409 aa for all *E. fawcettii* predicted genes. The mean GC content was 55.82% and the mean cysteine content was 3.4%, compared to 54.69% and 1.2%, respectively for all predicted genes of *E. fawcettii*. The high proportion (44.2%) of gene-sparse genes among prioritised CE’s was expected, as CE’s which were not classified as gene-dense were favoured during the prioritisation process, however high proportions of gene-sparse genes were also observed among the SP’s and CE’s (Table 4). Specifically, 26.8% of all *E. fawcettii* predicted genes were classed as gene-sparse, 26.3% as gene-dense and the remaining 46.9% classed as neither. In comparison, 31.9% of SP’s and 35.7% of CE’s were classed as gene-sparse and only 15.9% and 16.25%, respectively, were classed as gene-dense, indicating a preference for gene-sparse locations by proteins likely secreted by the pathogen. PolyAA repeat-containing proteins were not overrepresented among the prioritised CE’s, two were found to contain five consecutive Ala residues and one other contained five consecutive Arg residues. Additionally, no CE’s were found to contain SSR’s suggesting that diversity of *E. fawcettii* effector sequences is not being generated through an increased mutational rate related to short repetitive sequences. Furthermore, the prioritised CE’s were found scattered throughout the genome over 34 of the 141 gene-containing contigs and did not appear to cluster together. While AT-rich regions were not taken into consideration during the prioritisation of *E. fawcettii* CE’s, due to a low AT-rich coverage of 1.03%, it should be noted that higher proportions of SP’s and CE’s were found among genes on the edge of a contig and those within 2 Kb of an AT-rich region than expected. Out of the 252 genes found at the edge of a contig, 36 (14.2%) were SP’s and 11 (4.3%) were CE’s, compared to 12.7% and 2.0%, respectively, out of all *E. fawcettii* proteins. Similarly, of the 57 genes found within 2 Kb of an AT-rich region, 12 (21.1%) were SP’s and four (7.0%) were CE’s (S7). This suggests that genomic regions near contig breaks, such as sequences of low complexity or regions under-represented by short read sequencing technology, and AT-rich regions may be indicators within the *E. fawcettii* genome of nearby SP’s and effector genes. Interestingly, SP’s and CE’s were not overrepresented among genes found within 2 Kb of a predicted TE region, of the 120 genes found in these regions 12 (10%) were SP’s and 2 (1.7%) were CE’s, both slightly less than their proportions across the whole genome. This suggested while potential effector genes are more likely to be found near AT-rich regions, a nearby predictable TE region was not necessary. Thus, *E. fawcettii*, a necrotrophic pathogen not considered at first thought to utilise protein effectors to increase virulence, shows a subtle, yet intriguing, pattern of SP’s and CE’s near AT-rich regions, at contig edges and in more gene-sparse locations. This potentially points towards a set of virulence-related genes being maintained in specific genomic locations and therefore suggesting their potential significance.

**Figure 5.**
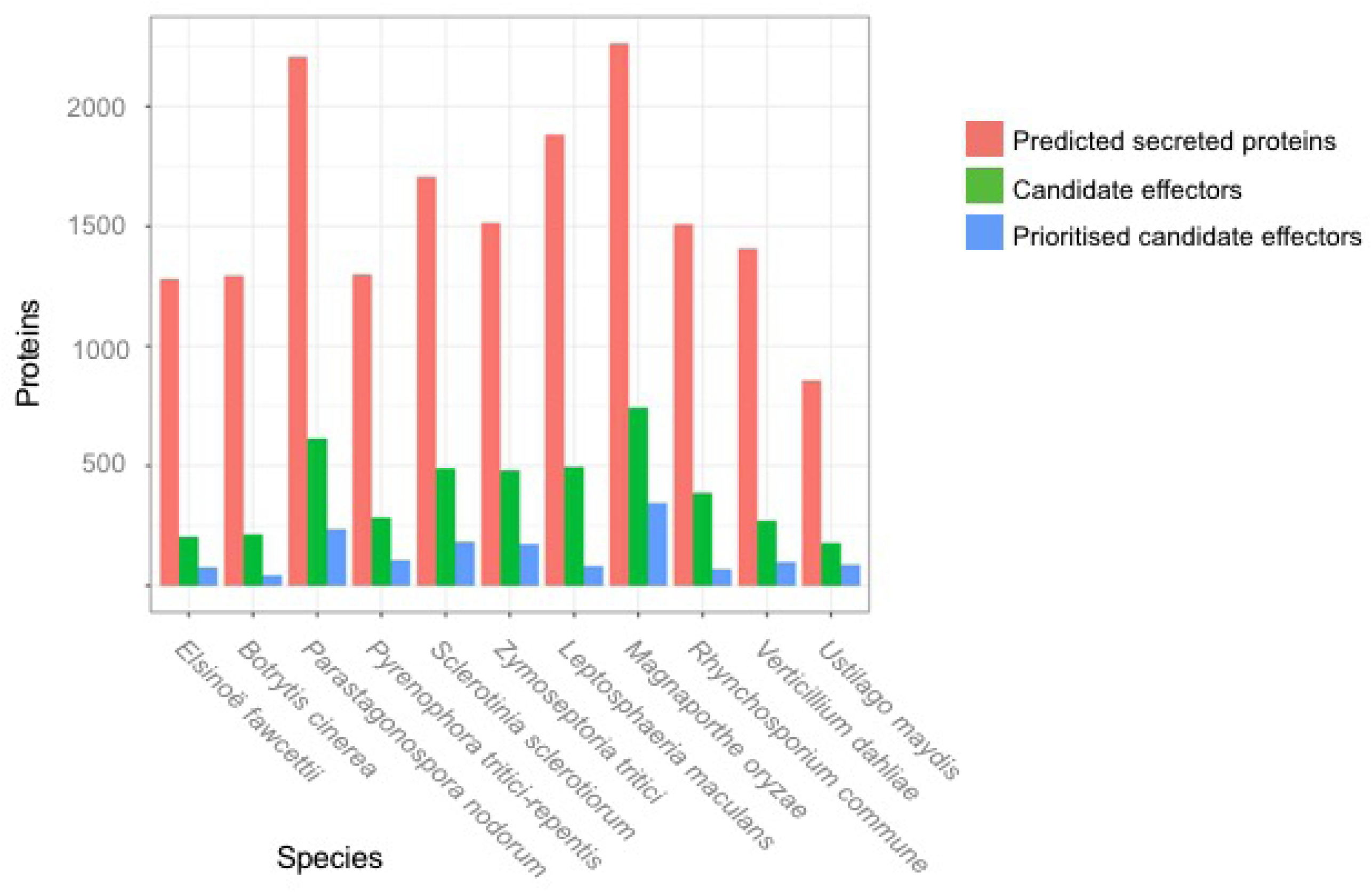
Comparison of numbers of secreted proteins, candidate effectors and prioritised candidate effectors among 11 fungal pathogens. Secreted proteins and candidate effectors were predicted using the pipeline in Figure 3. Prioritised candidate effectors were determined using features shown in Figure 4.

**Table 4.** Gene density classification of *Elsinoë fawcettii* predicted proteins

| Classification | All predicted proteins | Secreted proteins | Candidate effectors | Prioritised candidate effectors |
| --- | --- | --- | --- | --- |
| Gene-sparse | 26.8% | 31.9% | 35.7% | 44.2% |
| Gene-dense | 26.3% | 15.9% | 16.3% | 1.3% |
| Neither | 46.9% | 52.1% | 48.3% | 54.5% |

While analysing proteins using the features mentioned above can shortlist CE’s, awareness of limitations should be considered. For example, only prioritising CE’s which are unique to a species, or obtain the same orthoMCL hit as a known effector, limits the identification of novel effectors which may be utilised by multiple species. Hence, a blast search of *E. fawcettii* CE’s against CE’s of the 10 other fungal pathogens was conducted and indicated 12 (5.9%) *E. fawcettii* CE’s had >70% similarity to at least one candidate effector of another species (S7). Four of these 12 proteins were prioritised CE’s, one of which had 72.9% similarity to MoCDIP1 (*M. oryzae*), a known effector which is expressed in planta and induces host cell death [49], thus highlighting this CE for further investigation.

### Prediction and prioritisation of cell wall degrading enzymes

Further potential pathogenicity-related genes of *E. fawcettii* which deserve attention include CWDE’s. The *E. fawcettii* proteome showed 378 (3.75%) predicted CAZymes (S8), comparable to the proportion of CAZymes seen in the other 10 pathogen genomes, which ranged from 2.8% (*S. sclerotiorum*) to 4.3% (*V. dahliae*) (S2). Of the total *E. fawcettii* CAZymes, 203 (53.7%) were also predicted as secreted, highlighting numerous potential CWDE’s secreted by the pathogen and targeted for interaction with host carbohydrates. It would be beneficial to compare these potential CWDE’s with transcriptomic data once available, however, currently they can be cross-referenced against the Pfam database.

Analysis of the 203 potential CWDE’s revealed frequently appearing Pfam hits to pectate lyase and pectinesterase (19 hits), the glycosyl hydrolases family 28 of pectin-degrading polygalacturonases (11 hits) and the glycosyl hydrolases family 43 of hemicellulose-degrading beta-xylosidases (10 hits). Hemicellulose- and pectin-degrading enzymes target plant cell wall components including xyloglucans and pectin’s, respectively [68], both found in high proportions in the primary cell wall, potentially revealing an arsenal of CWDE’s of *E. fawcettii* which are targeted towards young plant tissues. Polygalacturonases break bonds between polygalacturonic acid residues, thereby degrading pectin, while beta-xylosidases hydrolyse xylan, a hemicellulose component of the cell wall. It is possible that the CWDE’s of *E. fawcettii* have the ability to degrade components of a growing cell wall, however as the host cell wall matures, the *E. fawcettii* CWDE repertoire becomes less effective, perhaps explaining why only young plant tissues are susceptible to citrus scab. The 203 potential CWDE’s were also cross-referenced against PHI-base, resulting in the prioritisation of 21 proteins which had similarity to known virulence factors of plant pathogens (Table 5, S8), thus highlighting candidate virulence genes of *E. fawcettii* for future experimental investigation. Among these 21 proteins were 14 predicted pectin-degrading enzymes, including two with similarity to polygalacturonase genes, specifically *pg1* (53.7%) and *pgx6* (66.4%) of *Fusarium oxysporum* which have been shown to reduce pathogen virulence when both are mutated simultaneously [74]; two showed similarity (61.6% and 41.8%) to the *PecA* polygalacturonase gene of *Aspergillus flavus*, a CWDE which primarily degrades pectin, and has been shown to improve pathogen invasion and increase spread during infection [73]; one with similarity to the pectin methylesterase *Bcpme1* gene of *B. cinerea* [78]; four with similarity (45.7% - 63.5%) to *PelA* and *PelD*, two pectate lyase virulence factors of *Nectria haematococca* [75]; and a further five obtained a pectate lyase Pfam hit, of which four showed similarity (40.3% - 53.5%) to the *Pnl1* pectin lyase gene of citrus pathogen *Penicillium digitatum* [76] and one with 58.4% similarity to *PelB* pectate lyase B gene of *Colletotrichum gloeosporioides*, seen to affect virulence on avocado [77]. A further five prioritised candidate CWDE’s, classed as hemicellulose-degrading enzymes, showed similarity (46.7% - 61.6%) to the endo-1,4-beta-xylanases (glycosyl hydrolase families 10 and 11) of *M. oryzae*, the knockdown of which is seen to reduce pathogenicity [80]. The remaining two prioritised CWDE’s, classed as cellulose-degrading enzymes, showed 51.9% and 52.9% similarity to the *Glu1* glucanase gene, a known virulence factor of wheat pathogen *Pyrenophora tritici-repentis* [79]. The similarities seen between these predicted secreted CAZymes and known virulence factors provides a collection of likely CWDE’s of *E. fawcettii* for future investigation. Unlike SP’s or CE’s, predicted CWDE’s of *E. fawcettii* were not overrepresented among genes found at the contig edge or within 2 Kb of an AT-rich region (S8). There was some crossover between CE’s and CWDE’s, with five *E. fawcettii* proteins being labelled as both prioritised CE’s and prioritised CWDE’s, thus providing some CE’s with potential carbohydrate-interacting functions.

**Table 5.**
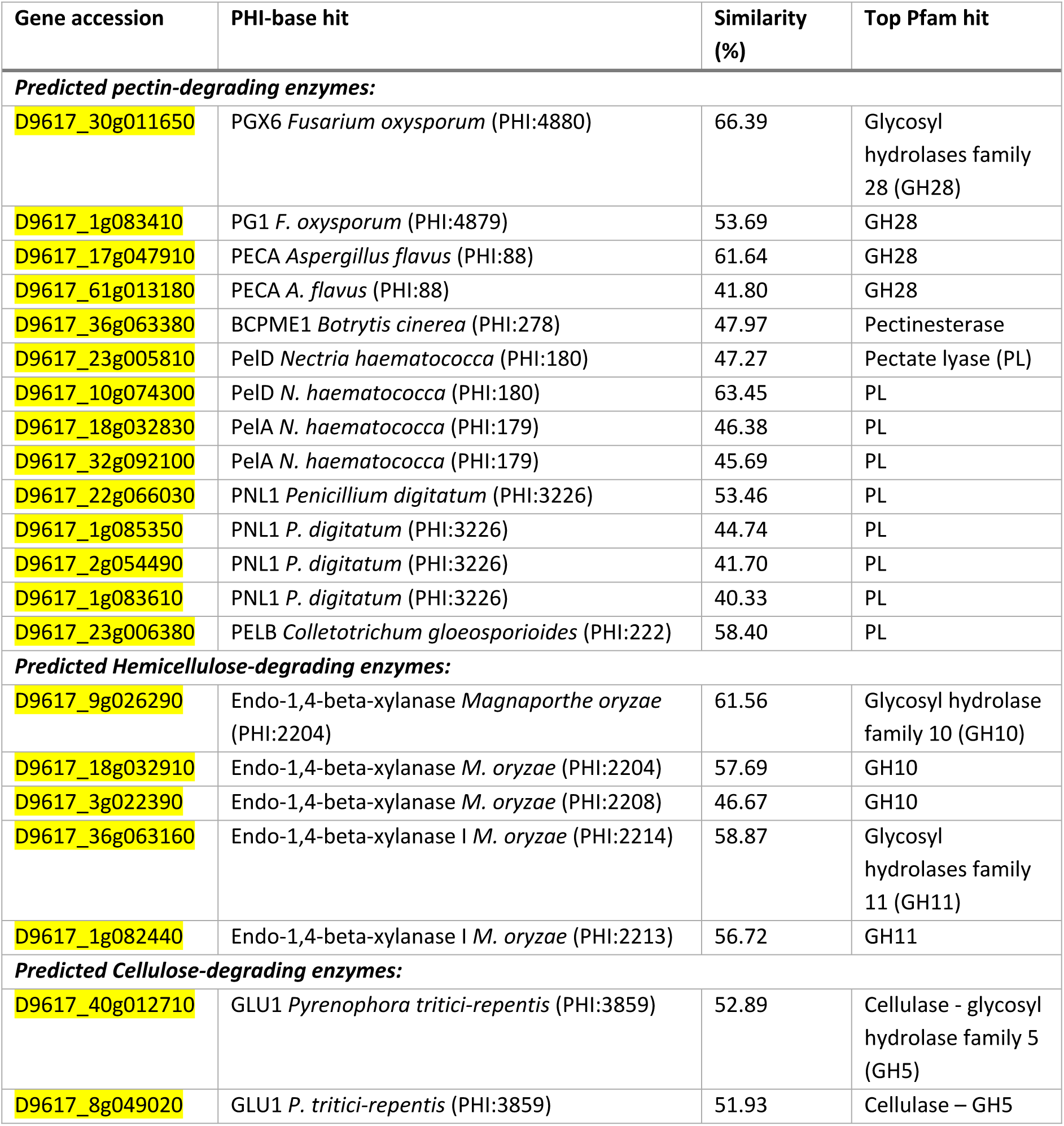
Predicted function of prioritised candidate cell wall degrading enzymes of *Elsinoë fawcettii*

### Prediction of secondary metabolite clusters

Much research surrounding *E. fawcettii* has focused on the SM elsinochrome, which contributes to the formation of necrotic lesions [25–28]. Analysis of the *E. fawcettii* genome assembly enabled the prediction of further genes potentially involved in the elsinochrome gene cluster than previously described, as well as the prediction of additional SM clusters throughout the assembly. In total, there were 22 predicted SM clusters, involving 404 (4.0%) genes (Table 6, S9). Comparing this to the results of the 10 comparative species showed that the number of predicted SM clusters varies widely among the pathogens, from 13 clusters (*U. maydis*) to 53 clusters (*M. oryzae*) (Figure 6). This wide variety among fungal species, in particular an overrepresentation of SM clusters among hemibiotrophs and necrotrophs has been seen before [154]. From the comparative analysis, it appears *E. fawcettii* has a lighter dependence upon the variety of secondary metabolite clusters compared to the other necrotrophs and hemibiotrophs, particularly for T1PKS clusters. Blast analysis of the previously determined *E. fawcettii* elsinochrome cluster [27] against the *E. fawcetti* proteome indicated high similarities in amino acid sequence for six genes of the predicted Type I Polyketide synthase (T1PKS) SM cluster 1 (S9). Specifically, the predicted core biosynthetic gene of cluster 1 (accession D9617_1g081920) showed 98.6% similarity to the *E. fawcettii* polyketide synthase (*EfPKS1*) gene (accession ABU63483.1). An additional predicted biosynthetic gene (accession D9617_1g081900) had 99.6% similarity to the *E. fawcettii* ESC reductase (*RDT1*) gene (accession ABZ01830) and the predicted transport-related gene (accession D9617_1g081940) showed 70.3% similarity to the *E. fawcettii* ECT1 transporter (*ECT1*) gene (accession ABZ82008). Additional genes within the *E. fawcettii* SM cluster 1 obtained hits to the *E. fawcettii* elsinochrome cluster [27], specifically D9617_1g081930, D9617_1g081910 and D9617_1g081890 had high (97.4% - 100%) similarity to *PRF1* prefoldin protein subunit 3 (accession ABZ01833.1), *TSF1* transcription factor (accession ABZ01831.1) and *EfHP1* coding a hypothetical protein (accession ABZ82009.1). Hence, SM cluster 1 contains the two genes, *EfPKS1* and *TSF1*, which have been shown to be essential in elsinochrome production, as well as four genes (*RDT1*, *PRF1*, *ECT1* and *EfHP1*) also thought to be involved in elsinochrome biosynthesis [26, 27]. SM cluster 1 appears to lack four genes, being *OXR1*, *EfHP2*, *EfHP3* and *EfHP4*, which have all been reported to code for hypothetical proteins and not thought to be involved in biosynthesis [27]. However, to further investigate these omissions, BLAST analysis querying the nucleotide sequences of the elsinochrome cluster [27] against the contigs of the *E. fawcettii* genome assembly indicated regions with high similarities (99.3% - 99.7%) consistent with the location of predicted SM cluster 1 on contig 1. This suggests that these unnecessary nearby genes may have become slightly degraded in the *E. fawcettii* BRIP 53147a isolate and were therefore not recognised during gene prediction. The use of alternate gene model prediction programs between the studies may also be a contributing factor. These differences may be further investigated through future transcriptomics analyses of *E. fawcettii*. Interestingly, SM cluster 1 consisted of an additional nine genes to the elsinochrome cluster previously described [27], all of which lay in a cluster adjacent to *ECT1*. Several of these additional genes obtained Pfam hits such as the THUMP domain, peptidase M3, Apolipoprotein O, Gar1/Naf1 RNA binding region and Endonuclease/Exonuclease/phosphatase family, suggesting these additional neighbouring proteins may perform functions such as RNA binding and modification, peptide cleavage, lipid binding and intracellular signalling, thus providing further genes for future investigation into the elsinochrome biosynthesis pathway.

**Figure 6.**
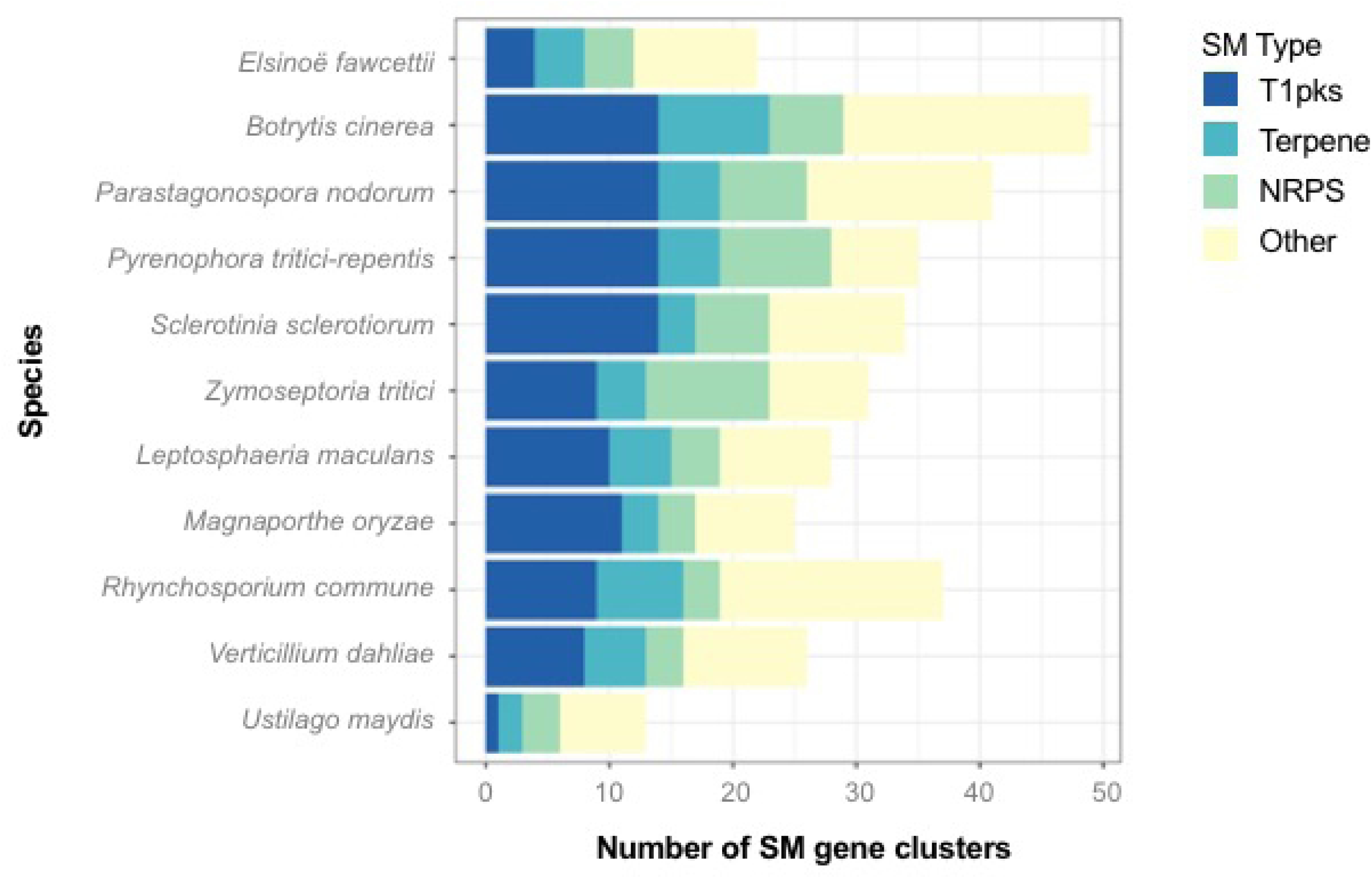
Comparison of numbers of predicted secondary metabolite gene clusters among 11 fungal species. Numbers of SM gene clusters, shown on the x axis, are divided into SM types; (I) Type I Polyketide synthase (T1PKS); (II) terpene; (III) non-ribosomal peptide synthetase (NRPS); and (IV) other, which contains all clusters identified by antiSMASH as either Type 3 Polyketide synthase (T3PKS), terpene-T1PKS, indole-T1PKS-NRPS, T1PKS-NRPS, indole-T1PKS, T1PKS-terpene-NRPS, indole, siderophore, lantipeptide, T3PKS-T1PKS or other.

**Table 6.**
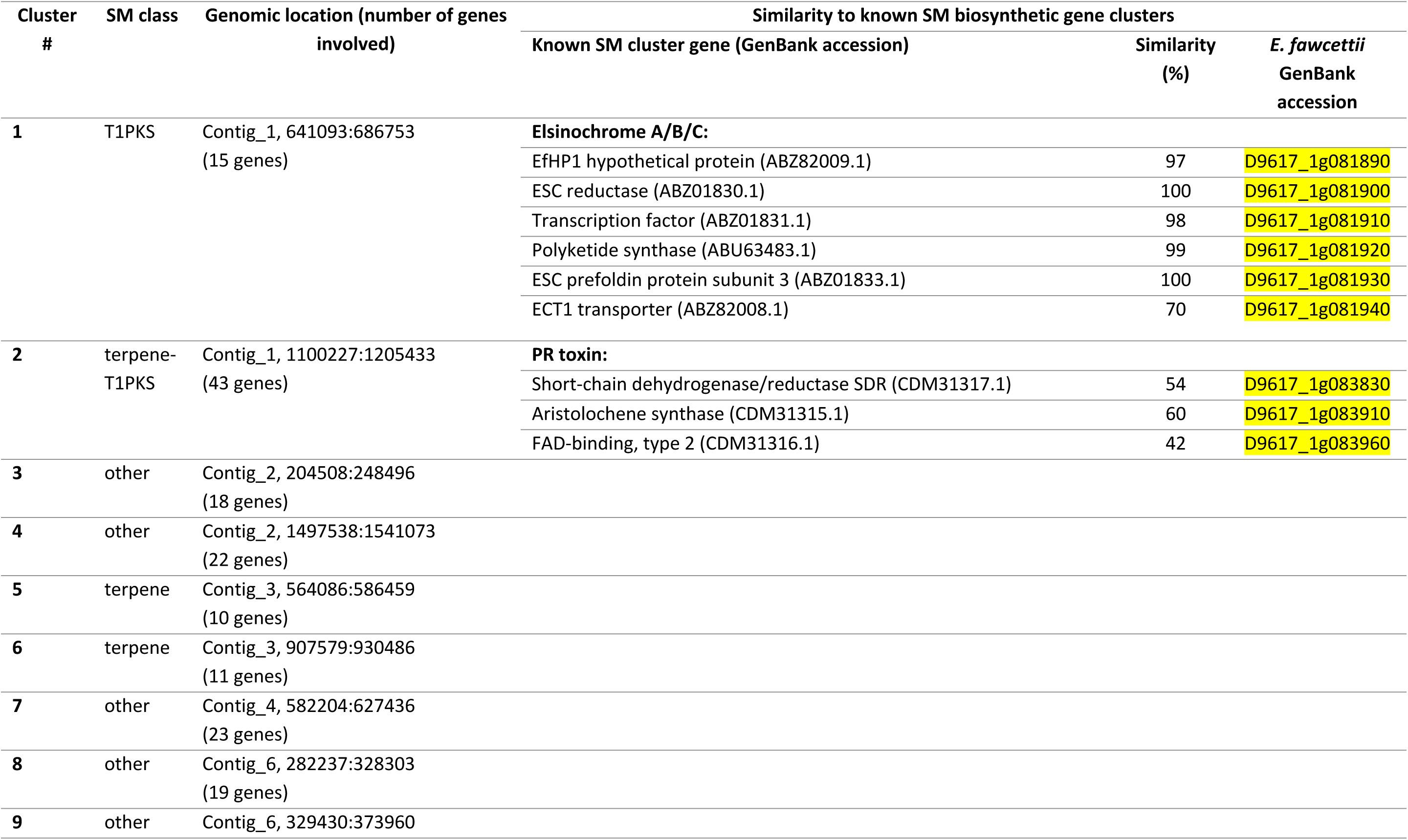
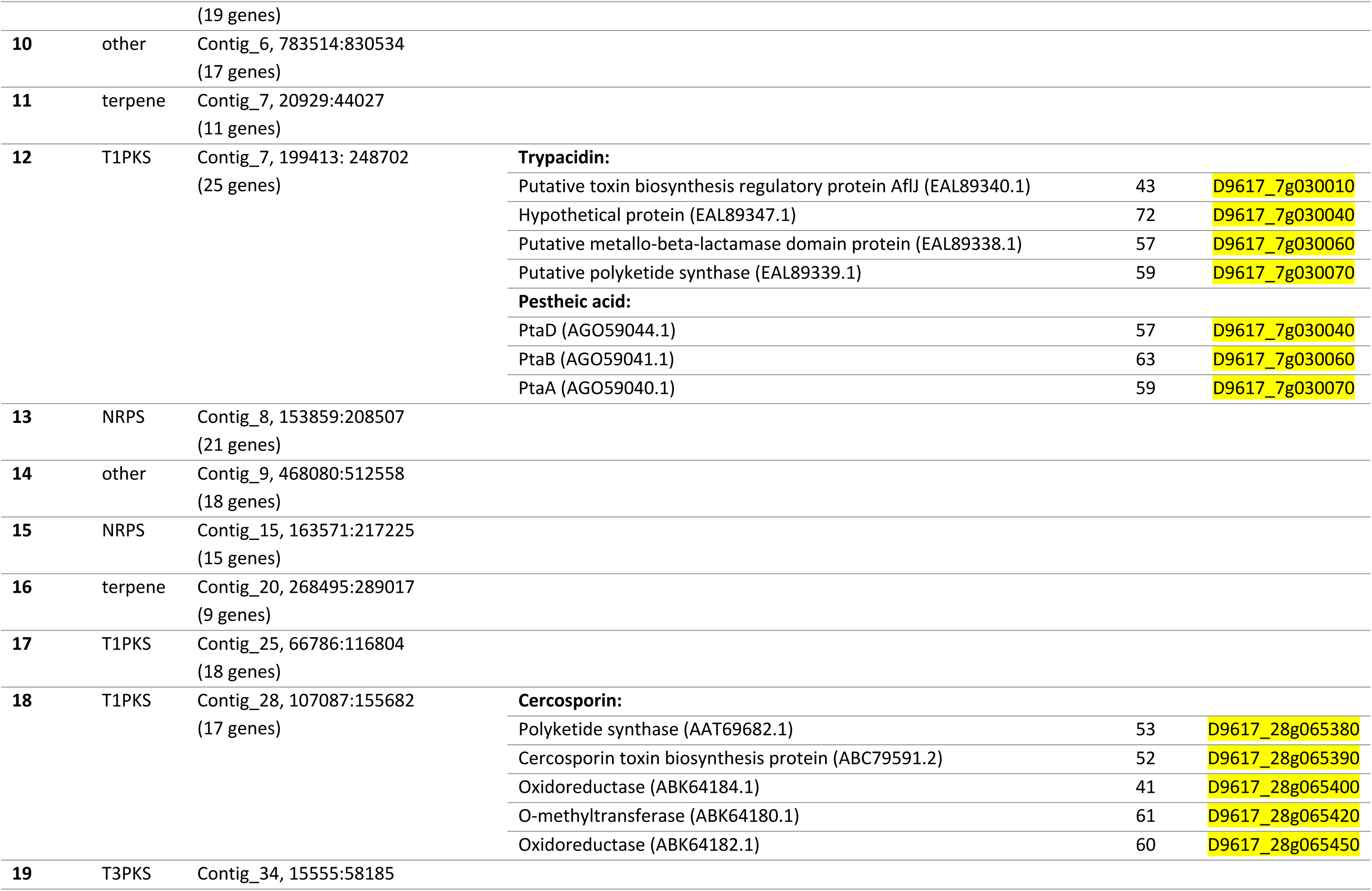
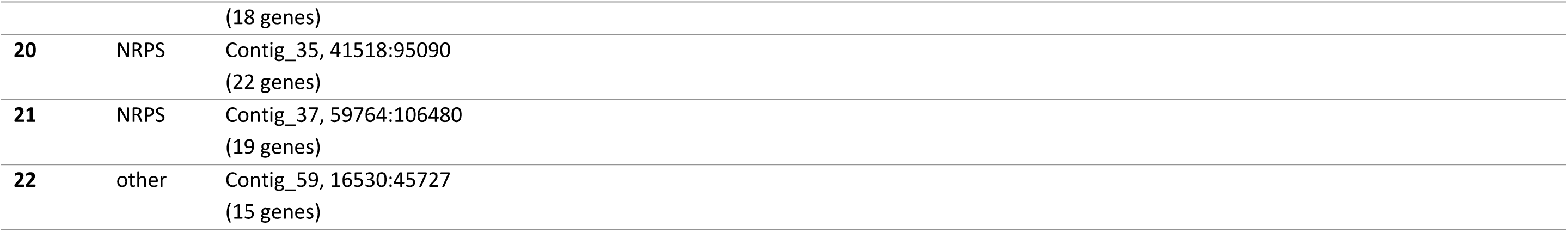
Predicted secondary metabolite (SM) gene clusters of *Elsinoë fawcettii*

An additional predicted SM cluster deserving of further investigation was SM cluster 2, a terpene-T1PKS, located 415,394 bp from the elsinochrome SM cluster 1 on contig 1. This cluster shows sequence similarity to three proteins within the PR toxin biosynthetic gene cluster, namely aristolochene synthase (accession CDM31315.1) with 60% similarity to D9617_1g083910, short-chain dehydrogenase/reductase (accession CDM31317.1) with 54% similarity D9617_1g083830 and the type 2 FAD-binding protein (accession CDM31316.1) with 42% similarity to D9617_1g083960. The PR toxin is produced by the saprobe *Penicillium roqueforti*, a known contaminant of silages [155], while the mechanisms of its likely role in plant degeneration are unknown [156], PR toxin is seen to induce necrosis in human intestinal epithelial cells and monocytic immune cells [157] and exhibits mutagenic activity towards rats [158]. Thus, indicating the potential production of a toxin by *E. fawcettii* with DNA-binding capabilities. Another predicted SM gene cluster of interest was the T1PKS SM cluster 12. Three genes of cluster 12 (D9617_7g030040, D9617_7g030060 and D9617_7g030070) showed similarity to multiple known biosynthetic genes clusters; including the pestheic acid biosynthetic gene cluster of *Pestalotiopsis fici* [159] thought to function as a plant growth regulator [160] and the Trypacidin biosynthetic gene cluster of *Aspergillus fumigatus*, which produces a SM toxic to human lung cells [161]. Lastly, SM cluster 18 is predicted to code for five proteins with sequence similarity to those of the cercosporin biosynthetic gene cluster of *Cercospora nicotianae* [162]. Specifically, D9617_28g065380 (53% similarity to polyketide synthase, accession AAT69682.1), D9617_28g065390 (52% similarity to cercosporin toxin biosynthesis protein, accession ABC79591.2), D9617_28g065400 (41% similarity to oxidoreductase, accession ABK64184.1), D9617_28g065420 (61% similarity to O-methyltransferase, accession ABK64180.1) and D9617_28g065450 (60% similarity to oxidoreductase, accession ABK64182.1). Cercosporin, similar to elsinochrome, is a fungal toxin which promotes the generation of reactive oxygen species in the presence of light, killing plant cells [163]. Cercosporin produced by *C. nicotianae* has been shown to cause necrotic lesions on tobacco leaves [164] and is also produced by the apple pathogen *Colletotrichum fioriniae* [165]. While it has been shown that elsinochrome production is important for full virulence by *E. fawcettii* [26, 27], biosynthesis of further SM’s, such as cluster 2, 12 or 18, may be beneficial to pathogenesis by potentially disrupting host plant signalling, causing additional necrosis or inhibiting competing microbes.

Analysis of the distances between predicted SM genes and TE’s indicated no TE’s were in the close vicinity of SM cluster 1 (elsinochrome), the closest TE to the edge of the cluster was 199,748 bp or 77 genes away. This lack of association was seen among all *E. fawcettii* predicted SM clusters, with seven clusters predicted on contigs without identified TE’s (S9). Of those clusters which did lie on contigs with TE’s, genes were an average distance of 236,556 bp away, suggesting recent activity of known TE’s was unlikely to be involved in the formation of *E. fawcettii* SM clusters. The closest AT-rich region to SM cluster 1 was a distance of 90,363 bp, while this was less than the mean distance (257,863 bp), this indication of potential TE degradation by RIP is still quite distant. In contrast to multiple SP’s and CE’s seen in the close vicinity of AT-rich regions, there were no genes from predicted SM clusters within 2 Kb of an AT-rich region, suggesting genes involved in SM production may benefit from residing in more stable genomic regions.

## Conclusion

The WGS sequencing, genome mining and comparative analyses conducted in this study illustrates the potential that exists within the genome of *E. fawcettii* for virulence factors such as protein effectors and CWDE’s. The identification of these potential pathogenicity-related genes is a first step in determining further mechanisms utilised by *E. fawcettii* in addition to elsinochrome production, thus enabling this pathogen to defeat plant immune strategies in a host-specific manner. This study provides predicted virulence genes for future experimental investigation of *E. fawcettii* pathogenesis pathways, as well as establishing a comprehensive genomic resource for use in future studies to determine improved methods of control and screening of this pathogen.

## Acknowledgments

We would like to thank members of the Centre for Crop Health at the University of Southern Queensland for their time and work. In particular, Lauren Huth and Katelynn Hadzi for providing laboratory and organisational support for the project, and Levente Kiss for providing guidance throughout this study.

## Supporting information captions

S1 Table. GenBank accessions for ITS and TEF1-α sequences included in the phylogenetic analysis with *Elsinoë fawcettii* isolate (BRIP 53147a).

S2 Table. Comparison of predicted gene classifications among *Elsinoë fawcettii* and 10 other species; Pfam hits, predicted CAZymes and core/acc/unique genes.

S3 Text. Sequence alignment of partial ITS and TEF1-α regions of *Elsinoë fawcettii* (BRIP 53147a) in comparison with other *E. fawcettii* isolates and closely related *Elsinoë* species.

S4 Table. Comparison of results of EffectorP predicted candidate effectors and alternate candidate effector search among 11 species.

S5 Table. Genomic and proteomic analyses of 11 species for use in known effector analysis and candidate effector prioritisation.

S6 Table. Comparison of numbers of predicted secreted proteins, candidate effectors and prioritised candidate effectors among 11 species.

S7 Table. Features and GenBank accessions of 203 *Elsinoë fawcettii* candidate effectors.

S8 Table. Features and GenBank accessions of 378 *Elsinoë fawcettii* predicted CAZymes.

S9 Table. Features and GenBank accessions of 404 *Elsinoë fawcettii* genes with predicted involvement in secondary metabolite clusters.

## References

1. Xin H, Huang F, Zhang T, Xu J, Hyde KD, Li H. Pathotypes and genetic diversity of Chinese collections of *Elsinoë fawcettii* causing citrus scab. Journal of Integrative Agriculture. 2014;13(6):1293–302.

2. Hyun J, Timmer L, Lee SC, Yun SH, Ko SW, Kim KS. Pathological characterization and molecular analysis of *Elsinoe* isolates causing scab diseases of citrus in Jeju Island in Korea. Plant disease. 2001;85(9):1013–7.

3. Hyun J, Yi S, MacKenzie S, Timmer L, Kim K, Kang S, et al. Pathotypes and genetic relationship of worldwide collections of *Elsinoë* spp. causing scab diseases of citrus. Phytopathology. 2009;99(6):721–8.

4. Tan M, Timmer L, Broadbent P, Priest M, Cain P. Differentiation by molecular analysis of *Elsinoë* spp. causing scab diseases of citrus and its epidemiological implications. Phytopathology. 1996;86(10):1039–44.

5. Bitancourt AA, Jenkins AE. Elsinoe fawcettii, the perfect stage of the Citrus scab fungus. Phytopathology. 1936;26(4):393–5.

6. Bitancourt AA, Jenkins AE. Sweet orange fruit scab caused by australis. Journal of Agricultural Research. 1937;54:0001–18.

7. Timmer L, Priest M, Broadbent P, Tan M. Morphological and pathological characterization of species of *Elsinoë* causing scab diseases of citrus. Phytopathology. 1996;86(10):1032–8.

8. Whiteside J. Biological characteristics of *Elsinoe fawcettii* pertaining to the epidemiology of sour orange scab. Phytopathology. 1975;65(10):1170–5.

9. Whiteside J. Pathogenicity of two biotypes of *Elsinoë fawcettii* to sweet orange and some other cultivars. Phytopathology. 1978;68(1).

10. Wang LY, Liao HL, Bau HJ, Chung KR. Characterization of pathogenic variants of *Elsinoë fawcettii* of citrus implies the presence of new pathotypes and cryptic species in Florida. Canadian Journal of Plant Pathology. 2009;31(1):28–37.

11. Queensland Government. Department of Agriculture and Fisheries. DAF Biological Collections [cited 2019 May 10]. Available from: http://collections.daff.qld.gov.au/web/home.html

12. dos Santos RF, Spósito MB, Ayres MR, Sosnowski MR. Phylogeny, morphology and pathogenicity of *Elsinoë ampelina*, the causal agent of grapevine anthracnose in Brazil and Australia. Journal of Phytopathology. 2018;166(3):187–98.

13. Ash G, Stodart B, Hyun JW. Black scab of jojoba (*Simmondsia chinensis*) in Australia caused by a putative new pathotype of *Elsinoë australis*. Plant disease. 2012;96(5):629–34.

14. Miles AK, Tan YP, Shivas RG, Drenth A. Novel pathotypes of Elsinoë australis associated with Citrus australasica and Simmondsia chinensis in Australia. Tropical Plant Pathology. 2015;40(1):26–34.

15. Kokoa P. Review of sweet potato diseases in PNG. Food Security for Papua New Guinea Proceedings of the Papua New Guinea Food and Nutrition 2000 Conference ACIAR Proceedings; 2001.

16. Mau YS. Resistance response of fifteen sweet potato genotypes to scab disease (*Sphaceloma batatas*) in two growing sites in East Nusa Tenggara, Indonesia. Tropical Drylands. 2018;2(1):5–11.

17. Scheper R, Wood P, Fisher B. Isolation, spore production and Koch’s postulates of Elsinoe pyri. NZ Plant Prot. 2013;66:308–16.

18. Fan X, Barreto R, Groenewald J, Bezerra J, Pereira O, Cheewangkoon R, et al. Phylogeny and taxonomy of the scab and spot anthracnose fungus *Elsinoë* (Myriangiales, Dothideomycetes). Studies in mycology. 2017;87:1–41.

19. Timmer LW, Garnsey SM, Graham JH. Compendium of Citrus Diseases. 2nd ed. The American Phytopathological Society. 2000.

20. Paudyal DP, Hyun JW. Physical changes in satsuma mandarin leaf after infection of *Elsinoë fawcettii* causing citrus scab disease. The plant pathology journal. 2015;31(4):421.

21. Agostini J, Bushong P, Bhatia A, Timmer L. Influence of environmental factors on severity of citrus scab and melanose. Plant disease. 2003;87(9):1102–6.

22. Chung KR. *Elsinoë fawcettii and Elsinoë australis*: the fungal pathogens causing citrus scab. Molecular plant pathology. 2011;12(2):123–35.

23. Weiss U, Ziffer H, Batterham T, Blumer M, Hackeng W, Copier H, et al. Pigments of *Elsinoë* species: Pigment production by *Elsinoë* species; isolation of pure elsinochromes A, B, and C. Canadian journal of microbiology. 1965;11(1):57–66.

24. Weiss U, Merlini L, Nasini G. Naturally occurring perylenequinones. Progress in the Chemistry of Organic Natural Products: Springer; 1987;52:1–71.

25. Liao HL, Chung KR. Cellular toxicity of elsinochrome phytotoxins produced by the pathogenic fungus, *Elsinoë fawcettii* causing citrus scab. New Phytologist. 2008;177(1):239–50.

26. Liao HL, Chung KR. Genetic dissection defines the roles of elsinochrome phytotoxin for fungal pathogenesis and conidiation of the citrus pathogen *Elsinoë fawcettii*. Molecular plant-microbe interactions. 2008;21(4):469–79.

27. Chung KR, Liao HL. Determination of a transcriptional regulator-like gene involved in biosynthesis of elsinochrome phytotoxin by the citrus scab fungus, Elsinoë fawcettii. Microbiology. 2008;154(11):3556–66.

28. Wang LY, Bau HJ, Chung KR. Accumulation of Elsinochrome phytotoxin does not correlate with fungal virulence among *Elsinoë fawcettii* isolates in Florida. Journal of phytopathology. 2009;157(10):602–8.

29. Hogenhout SA, Van Der Hoorn RAL, Terauchi R, Kamoun S. Emerging concepts in effector biology of plant-associated organisms. Molecular Plant-Microbe Interactions. 2009;22(2):115–22. doi: 10.1094/MPMI-22-2-0115.

30. Kamoun S. A catalogue of the effector secretome of plant pathogenic oomycetes. Annu Rev Phytopathol. 2006;44:41–60.

31. Bolton MD, Thomma BPHJ, Nelson BD. Sclerotinia sclerotiorum (Lib.) de Bary: biology and molecular traits of a cosmopolitan pathogen. Oxford, UK: Blackwell Science Ltd; 2006. p. 1–16.

32. Lyu X, Shen C, Fu Y, Xie J, Jiang D, Li G, et al. A small secreted virulence-related protein is essential for the necrotrophic interactions of *Sclerotinia sclerotiorum* with its host plants. PLoS Pathogens. 2016;12(2):e1005435. doi: 10.1371/journal.ppat.1005435.

33. Yu Y, Xiao J, Zhu W, Yang Y, Mei J, Bi C, et al. Ss-Rhs1, a secretory Rhs repeat-containing protein, is required for the virulence of *Sclerotinia sclerotiorum*. Molecular Plant Pathology. 2017;18(8):1052–61. doi: 10.1111/mpp.12459.

34. Friesen TL, Faris JD, Solomon PS, Oliver RP. Host-specific toxins: effectors of necrotrophic pathogenicity. Oxford, UK: Blackwell Publishing Ltd; 2008. p. 1421-8.

35. Rodriguez-Moreno L, Ebert MK, Bolton MD, Thomma BPHJ. Tools of the crook-infection strategies of fungal plant pathogens. Plant Journal. 2018;93(4):664–74. doi: 10.1111/tpj.13810.

36. Wang X, Jiang N, Liu J, Liu W, Wang G-L. The role of effectors and host immunity in plant–necrotrophic fungal interactions. Taylor & Francis; 2014. p. 722–32.

37. Lo Presti L, Lanver D, Schweizer G, Tanaka S, Liang L, Tollot M, et al. Fungal Effectors and Plant Susceptibility. Annual Review of Plant Biology. 2015;66(1):513–45. doi: 10.1146/annurev-arplant-043014-114623.

38. Stergiopoulos I, de Wit PJGM. Fungal Effector Proteins. Annual Review of Phytopathology. 2009;47(1):233–63. doi: 10.1146/annurev.phyto.112408.132637.

39. Manning VA, Pandelova I, Dhillon B, Wilhelm LJ, Goodwin SB, Berlin AM, et al. Comparative genomics of a plant-pathogenic fungus, Pyrenophora tritici-repentis, reveals transduplication and the impact of repeat elements on pathogenicity and population divergence. G3 (Bethesda, Md). 2013;3(1):41–63. doi: 10.1534/g3.112.004044.

40. Sperschneider J, Dodds PN, Gardiner DM, Manners JM, Singh KB, Taylor JM. Advances and challenges in computational prediction of effectors from plant pathogenic fungi. PLoS Pathogens. 2015;11(5):e1004806. doi: 10.1371/journal.ppat.1004806.

41. Martinez JP, Oesch NW, Ciuffetti LM. Characterization of the multiple-copy host-selective toxin gene, ToxB, in pathogenic and nonpathogenic isolates *of Pyrenaphora tritici-repentis*. Molecular Plant-Microbe Interactions. 2004;17(5):467–74. doi: 10.1094/MPMI.2004.17.5.467.

42. Friesen TL, Stukenbrock EH, Liu Z, Meinhardt S, Ling H, Faris JD, et al. Emergence of a new disease as a result of interspecific virulence gene transfer. Nature Genetics. 2006;38(8):953. doi: 10.1038/ng1839.

43. Syme RA, Hane JK, Friesen TL, Oliver RP. Resequencing and comparative genomics of *Stagonospora nodorum*: Sectional gene absence and effector discovery. G3: Genes, Genomes, Genetics. 2013;3(6):959–69. doi: 10.1534/g3.112.004994.

44. Chuma I, Isobe C, Hotta Y, Ibaragi K, Futamata N, Kusaba M, et al. Multiple translocation of the AVR-Pita effector gene among chromosomes of the rice blast fungus *Magnaporthe oryzae* and related species. PLoS Pathogens. 2011;7(7). doi: 10.1371/journal.ppat.1002147.

45. Ve T, Williams SJ, Catanzariti A-M, Rafiqi M, Rahman M, Ellis JG, et al. Structures of the flax-rust effector AvrM reveal insights into the molecular basis of plant-cell entry and effector-triggered immunity. Proceedings of the National Academy of Sciences of the United States. 2013;110(43):17594. doi: 10.1073/pnas.1307614110.

46. Kirsten S, Navarro-Quezada A, Penselin D, Wenzel C, Matern A, Leitner A, et al. Necrosis-inducing proteins of *Rhynchosporium commune*, effectors in quantitative disease resistance. Molecular Plant-Microbe Interactions. 2012;25(10):1314–25. doi: 10.1094/MPMI-03-12-0065-R.

47. Rouxel T, Grandaubert J, Hane JK, Hoede C, Van De Wouw AP, Couloux A, et al. Effector diversification within compartments of the *Leptosphaeria maculans* genome affected by Repeat-Induced Point mutations. Nature Communications. 2011;2(202):202. doi: 10.1038/ncomms1189.

48. Djamei A, Schipper K, Rabe F, Ghosh A, Vincon V, Kahnt J, et al. Metabolic priming by a secreted fungal effector. Nature. 2011;478(7369):395. doi: 10.1038/nature10454.

49. Chen S, Songkumarn P, Venu RC, Gowda M, Bellizzi M, Hu J, et al. Identification and characterization of in planta-expressed secreted effector proteins from *Magnaporthe oryzae* that induce cell death in rice. Molecular Plant-Microbe Interactions. 2013;26(2):191–202. doi: 10.1094/MPMI-05-12-0117-R.

50. Doehlemann G, van der Linde K, Assmann D, Schwammbach D, Hof A, Mohanty A, et al. Pep1, a secreted effector protein of *Ustilago maydis*, is required for successful invasion of plant cells. PLoS Pathogens. 2009;5(2). doi: 10.1371/journal.ppat.1000290.

51. Doehlemann G, Reissmann S, Aßmann D, Fleckenstein M, Kahmann R. Two linked genes encoding a secreted effector and a membrane protein are essential for *Ustilago maydis*-induced tumour formation. Molecular Microbiology. 2011;81(3):751–66. doi: 10.1111/j.1365-2958.2011.07728.x.

52. Mueller AN, Ziemann S, Treitschke S, Assmann D, Doehlemann G. Compatibility in the *Ustilago maydis*-maize interaction requires inhibition of host cysteine proteases by the fungal effector pit2. PLoS Pathogens. 2013;9(2). doi: 10.1371/journal.ppat.1003177.

53. Tanaka S, Brefort T, Neidig N, Djamei A, Kahnt J, Vermerris W, et al. A secreted *Ustilago maydis* effector promotes virulence by targeting anthocyanin biosynthesis in maize. eLife. 2014;2014(3). doi: 10.7554/eLife.01355.001.

54. Fudal I, Ross S, Gout L, Blaise F, Kuhn ML, Eckert MR, et al. Heterochromatin-like regions as ecological niches for avirulence genes in the *Leptosphaeria maculans* genome: Map-based cloning of AvrLm6. Molecular Plant-Microbe Interactions. 2007;20(4):459–70. doi: 10.1094/MPMI-20-4-0459.

55. Parlange F, Daverdin G, Fudal I, Kuhn ML, Balesdent MH, Blaise F, et al. *Leptosphaeria maculans* avirulence gene AvrLm4-7 confers a dual recognition specificity by the Rlm4 and Rlm7 resistance genes of oilseed rape, and circumvents Rlm4 -mediated recognition through a single amino acid change. Molecular Microbiology. 2009;71(4):851–63. doi: 10.1111/j.1365-2958.2008.06547.x.

56. Huang Y, Li Z, Evans N, Rouxel T, Fitt B, Balesdent M. Fitness Cost Associated with Loss of the AvrLm4 Avirulence Function in *Leptosphaeria maculans* (Phoma Stem Canker of Oilseed Rape). European Journal of Plant Pathology. 2006;114(1):77–89. doi: 10.1007/s10658-005-2643-4.

57. Saitoh H, Fujisawa S, Mitsuoka C, Ito A, Hirabuchi A, Ikeda K, et al. Large-scale gene disruption in *Magnaporthe oryzae* identifies MC69, a secreted protein required for infection by monocot and dicot fungal pathogens. PLoS Pathogens. 2012;8(5):e1002711. doi: 10.1371/journal.ppat.1002711.

58. Li W, Wang B, Wu J, Lu G, Hu Y, Zhang X, et al. The *Magnaporthe oryzae* avirulence gene AvrPiz-t encodes a predicted secreted protein that triggers the immunity in rice mediated by the blast resistance gene Piz-t. Molecular Plant-Microbe Interactions. 2009;22(4):411–20. doi: 10.1094/MPMI-22-4-0411.

59. Han L, Liu Z, Liu X, Qiu D. Purification, crystallization and preliminary X-ray diffraction analysis of the effector protein PevD1 from Verticillium dahliae. Acta Crystallographica Section F. 2012;68(7):802–5. doi: 10.1107/S1744309112020556.

60. Wang B, Yang X, Zeng H, Liu H, Zhou T, Tan B, et al. The purification and characterization of a novel hypersensitive-like response-inducing elicitor from *Verticillium dahliae* that induces resistance responses in tobacco. Applied Microbiology and Biotechnology. 2012;93(1):191–201. doi: 10.1007/s00253-011-3405-1.

61. Zhou R, Zhu T, Han L, Liu M, Xu M, Liu Y, et al. The asparagine-rich protein NRP interacts with the *Verticillium* effector PevD1 and regulates the subcellular localization of cryptochrome 2. Journal of Experimental Botany. 2017;68(13):3427–40. doi: 10.1093/jxb/erx192.

62. Schouten A, Van Baarlen P, Van Kan JAL. Phytotoxic Nep1-like proteins from the necrotrophic fungus *Botrytis cinerea* associate with membranes and the nucleus of plant cells. New Phytologist. 2008;177(2):493. doi: 10.1111/j.1469-8137.2007.02274.x.

63. Cuesta Arenas Y, Kalkman ERIC, Schouten A, Dieho M, Vredenbregt P, Uwumukiza B, et al. Functional analysis and mode of action of phytotoxic Nep1-like proteins of *Botrytis cinerea*. Physiological and Molecular Plant Pathology. 2010;74(5):376–86. doi: 10.1016/j.pmpp.2010.06.003.

64. Liu ZH, Faris JD, Meinhardt SW, Ali S, Rasmussen JB, Friesen TL. Genetic and physical mapping of a gene conditioning sensitivity in wheat to a partially purified host-selective toxin produced by *Stagonospora nodorum*. Phytopathology. 2004;94(10):1056–60. doi: 10.1094/PHYTO.2004.94.10.1056.

65. Martinez JP, Ottum SA, Ali S, Francl LJ, Ciuffetti LM. Characterization of the ToxB gene from *Pyrenophora tritici-repentis*. Molecular Plant-Microbe Interactions. 2001;14(5):675–7. doi: 10.1094/MPMI.2001.14.5.675.

66. Zhong Z, Marcel TC, Hartmann FE, Ma X, Plissonneau C, Zala M, et al. A small secreted protein in *Zymoseptoria tritici* is responsible for avirulence on wheat cultivars carrying the Stb6 resistance gene. New Phytologist. 2017;214(2):619–31. doi: 10.1111/nph.14434.

67. Raffaele S, Kamoun S. Genome evolution in filamentous plant pathogens: why bigger can be better. Nature Publishing Group; 2012. p. 417.

68. Kubicek CP, Starr TL, Glass NL. Plant cell wall degrading enzymes and their secretion in plant-pathogenic fungi. Annual Review of Phytopathology. 2014;52(1):427–51. doi: 10.1146/annurev-phyto-102313-045831.

69. King BC, Waxman KD, Nenni NV, Walker LP, Bergstrom GC, Gibson DM. Arsenal of plant cell wall degrading enzymes reflects host preference among plant pathogenic fungi. Biotechnology for Biofuels. 2011;4:4.

70. Cantarel BI, Coutinho PM, Rancurel C, Bernard T, Lombard V, Henrissat B. The Carbohydrate-Active EnZymes database (CAZy): An expert resource for glycogenomics. Nucleic Acids Research. 2009;37(1):D233–D8. doi: 10.1093/nar/gkn663.

71. Cosgrove DJ. Growth of the plant cell wall. Nature Publishing Group; 2005. p. 850.

72. Zuppini A, Navazio L, Sella L, Castiglioni C, Favaron F, Mariani P. An endopolygalacturonase from *Sclerotinia sclerotiorum* induces calcium-mediated signaling and programmed cell death in soybean cells. Molecular Plant-Microbe Interactions. 2005;18(8):849–55. doi: 10.1094/MPMI-18-0849.

73. Shieh MT, Brown RL, Whitehead MP, Cary JW, Cotty PJ, Cleveland TE, et al. Molecular genetic evidence for the involvement of a specific polygalacturonase, P2c, in the invasion and spread of *Aspergillus flavus* in cotton bolls. Applied and Environmental Microbiology. 1997;63(9):3548.

74. Bravo Ruiz G, Di Pietro A, Roncero MIG. Combined action of the major secreted exo- and endopolygalacturonases is required for full virulence of *Fusarium oxysporum*. Molecular Plant Pathology. 2016;17(3):339–53. doi: 10.1111/mpp.12283.

75. Rogers LM, Kim YK, Guo W, Gonzalez-Candelas L, Li D, Kolattukudy PE. Requirement for either a host- or pectin-induced pectate lyase for infection of Pisum sativum by Nectria hematococca. Proceedings of the National Academy of Sciences of the United States. 2000;97(17):9813. doi: 10.1073/pnas.160271497.

76. López-Pérez M, Ballester AR, González-Candelas L. Identification and functional analysis of *Penicillium digitatum* genes putatively involved in virulence towards citrus fruit. Molecular Plant Pathology. 2015;16(3):262–75. doi: 10.1111/mpp.12179.

77. Yakoby N, Beno-Moualem D, Keen NT, Dinoor A, Pines O, Prusky D. *Colletotrichum gloeosporioides* pelB is an important virulence factor in avocado fruit-fungus interaction. Molecular Plant-Microbe Interactions. 2001;14(8):988–95. doi: 10.1094/MPMI.2001.14.8.988.

78. Valette-Collet O, Cimerman A, Reignault P, Levis C, Boccara M. Disruption of *Botrytis cinerea* pectin methylesterase gene Bcpme1 reduces virulence on several host plants. Molecular Plant-Microbe Interactions. 2003;16(4):360–7. doi: 10.1094/MPMI.2003.16.4.360.

79. Fu H, Feng J, Aboukhaddour R, Cao T, Hwang SF, Strelkov SE. An exo-1,3-[beta]-glucanase GLU1 contributes to the virulence of the wheat tan spot pathogen *Pyrenophora tritici-repentis*. Fungal Biology. 2013;117(10):673.

80. Nguyen QB, Itoh K, Van Vu B, Tosa Y, Nakayashiki H. Simultaneous silencing of endo-β-1,4 xylanase genes reveals their roles in the virulence of *Magnaporthe oryzae*. Molecular Microbiology. 2011;81(4):1008–19. doi: 10.1111/j.1365-2958.2011.07746.x.

81. Brito N, Espino JJ, González C. The endo-β-1,4-xylanase Xyn11A is required for virulence in *Botrytis cinerea*. Molecular Plant-Microbe Interactions. 2006;19(1):25–32. doi: 10.1094/MPMI-19-0025.

82. Noda J, Brito N, Gonzalez C. The *Botrytis cinerea* xylanase Xyn11A contributes to virulence with its necrotizing activity, not with its catalytic activity. BMC Plant Biology. 2010;10:38.

83. Afgan E, Sloggett C, Goonasekera N, Makunin I, Benson D, Crowe M, et al. Genomics Virtual Laboratory: A Practical Bioinformatics Workbench for the Cloud. PLoS ONE. 2015;10(10):e0140829. doi: 10.1371/journal.pone.0140829.

84. Andrews S. FastQC: a quality control tool for high throughput sequence data. 2010.

85. Bolger AM, Lohse M, Usadel B. Trimmomatic: A flexible trimmer for Illumina sequence data. Bioinformatics. 2014;30(15):2114–20. doi: 10.1093/bioinformatics/btu170.

86. Zerbino DR, Birney E. Velvet: algorithms for de novo short read assembly using de Bruijn graphs. Genome Research. 2008;18(5):821–9. doi: 10.1101/gr.074492.107.

87. Gladman S. VelvetOptimiser. Victorian Bioinformatics Consortium, Clayton, Australia: http://bioinformaticsnetausoftwarevelvetoptimisershtml. 2012.

88. Bankevich A, Nurk S, Antipov D, Gurevich AA, Dvorkin M, Kulikov AS, et al. SPAdes: A New Genome Assembly Algorithm and Its Applications to Single-Cell Sequencing. Journal of Computational Biology. 2012;19(5):455–77. doi: 10.1089/cmb.2012.0021.

89. Langmead B, Salzberg SL. Fast gapped-read alignment with Bowtie 2. Nature methods. 2012;9(4):357.

90. The Picard Toolkit. http://broadinstitute.github.io/picard/ [cited 2018].

91. Thorvaldsdóttir H, Robinson JT, Mesirov JP. Integrative Genomics Viewer (IGV): high-performance genomics data visualization and exploration. Briefings in bioinformatics. 2013;14(2):178–92.

92. Simão FA, Waterhouse RM, Ioannidis P, Kriventseva EV, Zdobnov EM. BUSCO: Assessing genome assembly and annotation completeness with single-copy orthologs. Bioinformatics. 2015;31(19):3210–2. doi: 10.1093/bioinformatics/btv351.

93. Zdobnov EM, Tegenfeldt F, Kuznetsov D, Waterhouse RM, Simao FA, Ioannidis P, et al. OrthoDB v9.1: Cataloging evolutionary and functional annotations for animal, fungal, plant, archaeal, bacterial and viral orthologs. Nucleic Acids Research. 2017;45(1):D744–D9. doi: 10.1093/nar/gkw1119.

94. Testa AC, Oliver RP, Hane JK. OcculterCut: a comprehensive survey of AT-rich regions in fungal genomes. Genome biology and evolution. 2016;8(6):2044–64.

95. Lee T, Peace C, Jung S, Zheng P, Main D, Cho I. GenSAS - An online integrated genome sequence annotation pipeline. 2011. p. 1967–73.

96. Lomsadze A, Ter-Hovhannisyan V, Chernoff YO, Borodovsky M. Gene identification in novel eukaryotic genomes by self-training algorithm. Nucleic Acids Research. 2005;33(20):6494–506. doi: 10.1093/nar/gki937.

97. Smit AFA, Hubley R, Green P. RepeatMasker http://repeatmasker.org.

98. Beier S, Thiel T, Münch T, Scholz U, Mascher M. MISA-web: a web server for microsatellite prediction. Bioinformatics (Oxford, England). 2017;33(16):2583–5. doi: 10.1093/bioinformatics/btx198.

99. Altschul SF, Madden TL, Schäffer AA, Zhang J, Zhang Z, Miller W, et al. Gapped BLAST and PSI-BLAST: A new generation of protein database search programs. Nucleic Acids Research. 1997;25(17):3389–402. doi: 10.1093/nar/25.17.3389.

100. The Uniprot Consortium. UniProt: the universal protein knowledgebase. Nucleic acids research. 2018;46(5):2699.

101. Conesa A, Götz S, García-Gómez JM, Terol J, Talón M, Robles M. Blast2GO: A universal tool for annotation, visualization and analysis in functional genomics research. Bioinformatics. 2005;21(18):3674–6. doi: 10.1093/bioinformatics/bti610.

102. Johnson LS, Eddy SR, Portugaly E. Hidden Markov model speed heuristic and iterative HMM search procedure. BMC Bioinformatics. 2010;11(1). doi: 10.1186/1471-2105-11-431.

103. Finn RD, Coggill P, Eberhardt RY, Eddy SR, Mistry J, Mitchell AL, et al. The Pfam protein families database: Towards a more sustainable future. Nucleic Acids Research. 2016;44(1):D279–D85. doi: 10.1093/nar/gkv1344.

104. Quinlan AR. BEDTools: the Swiss-army tool for genome feature analysis. Current protocols in bioinformatics. 2014;47(1):11.2. 1-.2. 34.

105. Grant CE, Bailey TL, Noble WS. FIMO: Scanning for occurrences of a given motif. Bioinformatics. 2011;27(7):1017–8. doi: 10.1093/bioinformatics/btr064.

106. Bailey TL, Boden M, Buske FA, Frith M, Grant CE, Clementi L, et al. MEME Suite: Tools for motif discovery and searching. Nucleic Acids Research. 2009;37(2):W202–W8. doi: 10.1093/nar/gkp335.

107. Edgar RC. MUSCLE: Multiple sequence alignment with high accuracy and high throughput. Nucleic Acids Research. 2004;32(5):1792–7. doi: 10.1093/nar/gkh340.

108. Kumar S, Stecher G, Tamura K. MEGA7: Molecular Evolutionary Genetics Analysis Version 7.0 for Bigger Datasets. Molecular biology and evolution. 2016;33(7):1870–4. doi: 10.1093/molbev/msw054

109. Nei M. Molecular evolution and phylogenetics. Kumar S, editor. Oxford;: Oxford University Press; 2000.

110. Maddison WP, Maddison DR. Mesquite: a modular sysem for evolutionary analysis. Version 3.6 http://www.mesquiteproject.org 2018.

111. Kämper J, Kahmann R, Bölker M, Ma L-J, Brefort T, Saville BJ, et al. Insights from the genome of the biotrophic fungal plant pathogen Ustilago maydis. Nature. 2006;444(7115):97.

112. Rouxel T, Grandaubert J, Hane JK, Hoede C, Van De Wouw AP, Couloux A, et al. Effector diversification within compartments of the *Leptosphaeria maculans* genome affected by Repeat-Induced Point mutations. Nature Communications. 2011;2(202):202. doi: 10.1038/ncomms1189.

113. Dean RA, Talbot NJ, Ebbole DJ, Farman ML, Mitchell TK, Orbach MJ, et al. The genome sequence of the rice blast fungus *Magnaporthe grisea*. Nature. 2005;434(7036):980.

114. Penselin D, Münsterkötter M, Kirsten S, Felder M, Taudien S, Platzer M, et al. Comparative genomics to explore phylogenetic relationship, cryptic sexual potential and host specificity of *Rhynchosporium* species on grasses. BMC genomics. 2016;17(1):953.

115. Klosterman S, Subbarao K, Kang S, Veronese P, Gold S, Thomma B, et al. Comparative genomics of the plant vascular wilt pathogens, *Verticillium dahliae* and *Verticillium alboatrum*. Phytopathology. 2010;100(6):S64-S.

116. Van Kan JA, Stassen JH, Mosbach A, Van Der Lee TA, Faino L, Farmer AD, et al. A gapless genome sequence of the fungus *Botrytis cinerea*. Molecular plant pathology. 2017;18(1):75–89.

117. Hane JK, Lowe RG, Solomon PS, Tan K-C, Schoch CL, Spatafora JW, et al. Dothideomycete–plant interactions illuminated by genome sequencing and EST analysis of the wheat pathogen *Stagonospora nodorum*. The Plant Cell. 2007;19(11):3347–68.

118. Moolhuijzen P, See PT, Hane JK, Shi G, Liu Z, Oliver RP, et al. Comparative genomics of the wheat fungal pathogen *Pyrenophora tritici-repentis* reveals chromosomal variations and genome plasticity. BMC genomics. 2018;19(1):279.

119. Amselem J, Cuomo CA, van Kan JAL, Viaud M, Benito EP, Couloux A, et al. Genomic analysis of the necrotrophic fungal pathogens *Sclerotinia sclerotiorum* and *Botrytis cinerea*. PLoS Genetics. 2011;7(8):e1002230. doi: 10.1371/journal.pgen.1002230.

120. Plissonneau C, Hartmann FE, Croll D. Pangenome analyses of the wheat pathogen *Zymoseptoria tritici* reveal the structural basis of a highly plastic eukaryotic genome. BMC biology. 2018;16(1):5.

121. Sperschneider J, Dodds PN, Gardiner DM, Singh KB, Taylor JM. Improved prediction of fungal effector proteins from secretomes with EffectorP 2.0. Molecular Plant Pathology. 2018;19(9):2094–110. doi: 10.1111/mpp.12682.

122. Petersen TN, Brunak S, Heijne GV, Nielsen H. SignalP 4.0: discriminating signal peptides from transmembrane regions. Nature Methods. 2011;8(10):785. doi: 10.1038/nmeth.1701.

123. Käll L, Krogh A, Sonnhammer ELL. A Combined Transmembrane Topology and Signal Peptide Prediction Method. Journal of Molecular Biology. 2004;338(5):1027–36. doi: 10.1016/j.jmb.2004.03.016.

124. SoftBerry Inc. ProtComp v6 [cited 2018 July 19]. Available from: http://www.softberry.com/berry.phtml?topic=fdp.htm&no_menu=on.

125. Krogh A, Larsson B, Von Heijne G, Sonnhammer ELL. Predicting transmembrane protein topology with a hidden markov model: application to complete genomes. Journal of Molecular Biology. 2001;305(3):567–80. doi: 10.1006/jmbi.2000.4315.

126. Pierleoni A, Martelli P, Casadio R. PredGPI: A GPI-anchor predictor. BMC Bioinformatics. 2008;9(1). doi: 10.1186/1471-2105-9-392.

127. R Core Team. R: A language and environment for statistical computing. R Foundation for Statistical Computing, Vienna, Austria https://www.R-project.org/ 2018.

128. Blin K, Wolf T, Chevrette MG, Lu X, Schwalen CJ, Kautsar SA, et al. AntiSMASH 4.0 - improvements in chemistry prediction and gene cluster boundary identification. Nucleic Acids Research. 2017;45(1):W36–W41. doi: 10.1093/nar/gkx319.

129. Fischer S, Brunk BP, Chen F, Gao X, Harb OS, Iodice JB, et al. Using OrthoMCL to assign proteins to OrthoMCL-DB groups or to cluster proteomes into new ortholog groups. Current protocols in bioinformatics. 2011;35(1):6.12. 1–6.. 9.

130. Steiner L, Findeiß S, Lechner M, Marz M, Stadler Peter F, Prohaska Sonja J. Proteinortho: Detection of (Co-)orthologs in large-scale analysis. BMC Bioinformatics. 2011;12(1):124. doi: 10.1186/1471-2105-12-124.

131. Zhang H, Yohe T, Huang L, Entwistle S, Wu P, Yang Z, et al. DbCAN2: A meta server for automated carbohydrate-active enzyme annotation. Nucleic Acids Research. 2018;46(1):W95–W101. doi: 10.1093/nar/gky418.

132. Yin Y, Mao X, Yang J, Chen X, Mao F, Xu Y. DbCAN: A web resource for automated carbohydrate-active enzyme annotation. Nucleic Acids Research. 2012;40(1):W445–W51. doi: 10.1093/nar/gks479.

133. Buchfink B, Xie C, Huson D, H. Fast and sensitive protein alignment using DIAMOND. Nature Methods. 2014;12(1). doi: 10.1038/nmeth.3176.

134. Lombard V, Golaconda Ramulu H, Drula E, Coutinho PM, Henrissat B. The carbohydrate-active enzymes database (CAZy) in 2013. Nucleic Acids Research. 2014;42(1):D490–D5. doi: 10.1093/nar/gkt1178.

135. Busk PK, Pilgaard B, Lezyk MJ, Meyer AS, Lange L. Homology to peptide pattern for annotation of carbohydrate-active enzymes and prediction of function.(Report). BMC Bioinformatics. 2017;18(1). doi: 10.1186/s12859-017-1625-9.

136. Urban M, Cuzick A, Rutherford K, Irvine A, Pedro H, Pant R, et al. PHI-base: A new interface and further additions for the multi-species pathogen-host interactions database. Nucleic Acids Research. 2017;45(1):D604–D10. doi: 10.1093/nar/gkw1089.

137. Kis-Papo T, Weig AR, Riley R, Peršoh D, Salamov A, Sun H, et al. Genomic adaptations of the halophilic Dead Sea filamentous fungus *Eurotium rubrum*. Nature Communications. 2014;5(1). doi: 10.1038/ncomms4745.

138. Gazis R, Kuo A, Riley R, Labutti K, Lipzen A, Lin J, et al. The genome *of Xylona heveae* provides a window into fungal endophytism. Fungal Biology. 2016;120(1):26–42. doi: 10.1016/j.funbio.2015.10.002.

139. Rosienski MD, Lee MK, Yu JH, Kaspar CW, Gibbons JG. Genome sequence of the extremely acidophilic fungus Acidomyces richmondensis FRIK2901. Microbiology Resource Announcements. 2018;7(16). doi: 10.1128/MRA.01314-18.

140. Mohanta TK, Bae H. The diversity of fungal genome. BioMed Central Ltd.; 2015.

141. Bao W, Kojima K, Kohany O. Repbase Update, a database of repetitive elements in eukaryotic genomes. Mobile DNA. 2015;6(1). doi: 10.1186/s13100-015-0041-9.

142. Thomma BPHJ, Seidl MF, Shi-Kunne X, Cook DE, Bolton MD, van Kan JAL, et al. Mind the gap; seven reasons to close fragmented genome assemblies. Fungal Genetics and Biology. 2016;90:24–30. doi: 10.1016/j.fgb.2015.08.010.

143. Amselem J, Cuomo CA, van Kan JAL, Viaud M, Benito EP, Couloux A, et al. Genomic analysis of the necrotrophic fungal pathogens *Sclerotinia sclerotiorum* and *Botrytis cinerea*. PLoS Genetics. 2011;7(8):e1002230. doi: 10.1371/journal.pgen.1002230.

144. Cambareri EB, Jensen BC, Schabtach E, Selker EU. Repeat-induced G-C to A-T mutations in *Neurospora*. (glycine-cysteine to alanine-threonine). Science. 1989;244(4912):1571. doi: 10.1126/science.2544994.

145. Selker EU, Cambareri EB, Jensen BC, Haack KR. Rearrangement of duplicated DNA in specialized cells of *Neurospora*. Cell. 1987;51(5):741–52. doi: 10.1016/0092-8674(87)90097-3.

146. Selker EU. Premeiotic Instability of Repeated Sequences in Neurospora Crassa. Annual Review of Genetics. 1990;24(1):579–613. doi: 10.1146/annurev.ge.24.120190.003051.

147. Gladyshev E. Repeat-Induced Point Mutation (RIP) and Other Genome Defense Mechanisms in Fungi. Microbiology spectrum. 2017;5(4).

148. Braumann I, Berg M, Kempken F. Repeat induced point mutation in two asexual fungi, *Aspergillus niger* and *Penicillium chrysogenum*. Current Genetics. 2008;53(5):287–97. doi: 10.1007/s00294-008-0185-y.

149. Gout L, Fudal I, Kuhn ML, Blaise F, Eckert M, Cattolico L, et al. Lost in the middle of nowhere: the AvrLm1 avirulence gene of the Dothideomycete *Leptosphaeria maculans*. Molecular Microbiology. 2006;60(1):67–80. doi: 10.1111/j.1365-2958.2006.05076.x.

150. Fudal I, Ross S, Brun H, Besnard AL, Ermel M, Kuhn ML, et al. Repeat-Induced Point Mutation (RIP) as an alternative mechanism of evolution toward virulence in *Leptosphaeria maculans*. Molecular Plant-Microbe Interactions. 2009;22(8):932–41. doi: 10.1094/MPMI-22-8-0932.

151. Kloppholz S, Kuhn H, Requena N. A Secreted Fungal Effector of *Glomus intraradices* Promotes Symbiotic Biotrophy. Current Biology. 2011;21(14):1204–9. doi: 10.1016/j.cub.2011.06.044.

152. Mesarich CH, Bowen JK, Hamiaux C, Templeton MD. Repeat-containing protein effectors of plant-associated organisms. Frontiers in Plant Science. 2015;6. doi: 10.3389/fpls.2015.00872.

153. Liu T, Song T, Zhang X, Yuan H, Su L, Li W, et al. Unconventionally secreted effectors of two filamentous pathogens target plant salicylate biosynthesis. Nature Communications. 2014;5(1). doi: 10.1038/ncomms5686.

154. Zuccaro A, Lahrmann U, Langen G. Broad compatibility in fungal root symbioses. Current Opinion in Plant Biology. 2014;20:135–45. doi: 10.1016/j.pbi.2014.05.013.

155. Rasmussen R, Storm I, Rasmussen P, Smedsgaard J, Nielsen K. Multi-mycotoxin analysis of maize silage by LC-MS/MS. Analytical and Bioanalytical Chemistry. 2010;397(2):765–76. doi: 10.1007/s00216-010-3545-7.

156. Dubey MK, Aamir M, Kaushik MS, Khare S, Meena M, Singh S, et al. PR Toxin - Biosynthesis, Genetic Regulation, Toxicological Potential, Prevention and Control Measures: Overview and Challenges. Frontiers in Pharmacology. 2018;9. doi: 10.3389/fphar.2018.00288.

157. Hymery N, Puel O, Tadrist S, Canlet C, Le Scouarnec H, Coton E, et al. Effect of PR toxin on THP1 and Caco-2 cells: an in vitro study. World Mycotoxin Journal. 2017;10(4):375–86.

158. Polonelli L, Lauriola L, Morace G. Preliminary studies on the carcinogenic effects of *Penicillium roqueforti* toxin (PR toxin) on rats. Mycopathologia. 1982;78(2):125–7. doi: 10.1007/BF00442636.

159. Xu X, Liu L, Zhang F, Wang W, Li J, Guo L, et al. Identification of the First Diphenyl Ether Gene Cluster for Pestheic Acid Biosynthesis in Plant Endophyte *Pestalotiopsis fici*. ChemBioChem. 2014;15(2):284–92. doi: 10.1002/cbic.201300626.

160. Shimada A, Takahashi I, Kawano T, Kimura Y. Chloroisosulochrin, Chloroisosulochrin Dehydrate, and Pestheic Acid, Plant Growth Regulators, Produced by Pestalotiopsis theae. Zeitschrift fur Naturforschung - Section B Journal of Chemical Sciences. 2001;56(8):797–803. doi: 10.1515/znb-2001-0813.

161. Gauthier T, Wang X, Dos Santos JS, Fysikopoulos A, Tadrist S, Canlet C, et al. Trypacidin, a spore-borne toxin from *Aspergillus fumigatus*, is cytotoxic to lung cells. PLoS ONE. 2012;7(2):e29906. doi: 10.1371/journal.pone.0029906.

162. Chen H, Lee MH, Daub ME, Chung KR. Molecular analysis of the cercosporin biosynthetic gene cluster in *Cercospora nicotianae*. Molecular Microbiology. 2007;64(3):755–70. doi: 10.1111/j.1365-2958.2007.05689.x.

163. Daub ME, Hangarter RP. Light-induced production of singlet oxygen and superoxide by the fungal toxin, cercosporin. Plant Physiology. 1983;73(3):855–7.

164. Dekkers KL, You B-J, Gowda VS, Liao H-L, Lee M-H, Bau H-J, et al. The *Cercospora nicotianae* gene encoding dual O-methyltransferase and FAD-dependent monooxygenase domains mediates cercosporin toxin biosynthesis. Fungal Genetics and Biology. 2007;44(5):444–54. doi: 10.1016/j.fgb.2006.08.005.

165. de Jonge R, Ebert MK, Huitt-Roehl CR, Pal P, Suttle JC, Spanner RE, et al. Gene cluster conservation provides insight into cercosporin biosynthesis and extends production to the genus Colletotrichum. Proceedings of the National Academy of Sciences of the United States. 2018;115(24):E5459. doi: 10.1073/pnas.1712798115.

